# Simple and efficient modification of Golden Gate design standards and parts using oligo stitching

**DOI:** 10.1101/2022.02.10.479870

**Authors:** Jonas De Saeger, Mattias Vermeersch, Christophe Gaillochet, Thomas B. Jacobs

## Abstract

The assembly of DNA parts is a critical aspect of contemporary biological research. Gibson assembly and Golden Gate cloning are two popular options. Here, we explore the use of single stranded DNA oligos with Gibson assembly to augment Golden Gate cloning workflows in a process called “oligo stitching”. Our results show that oligo stitching can efficiently convert Golden Gate parts between different assembly standards and directly assemble incompatible Golden Gate parts without PCR amplification. Building on previous reports, we show that it can also be used to assemble *de novo* sequences. As a final application, we show that restriction enzyme recognition sites can be removed from plasmids and utilize the same concept to perform saturation mutagenesis. Given oligo stitching’s versatility and high efficiency, we expect that it will be a useful addition to the molecular biologist’s toolbox.

**Abstract Graphic:** 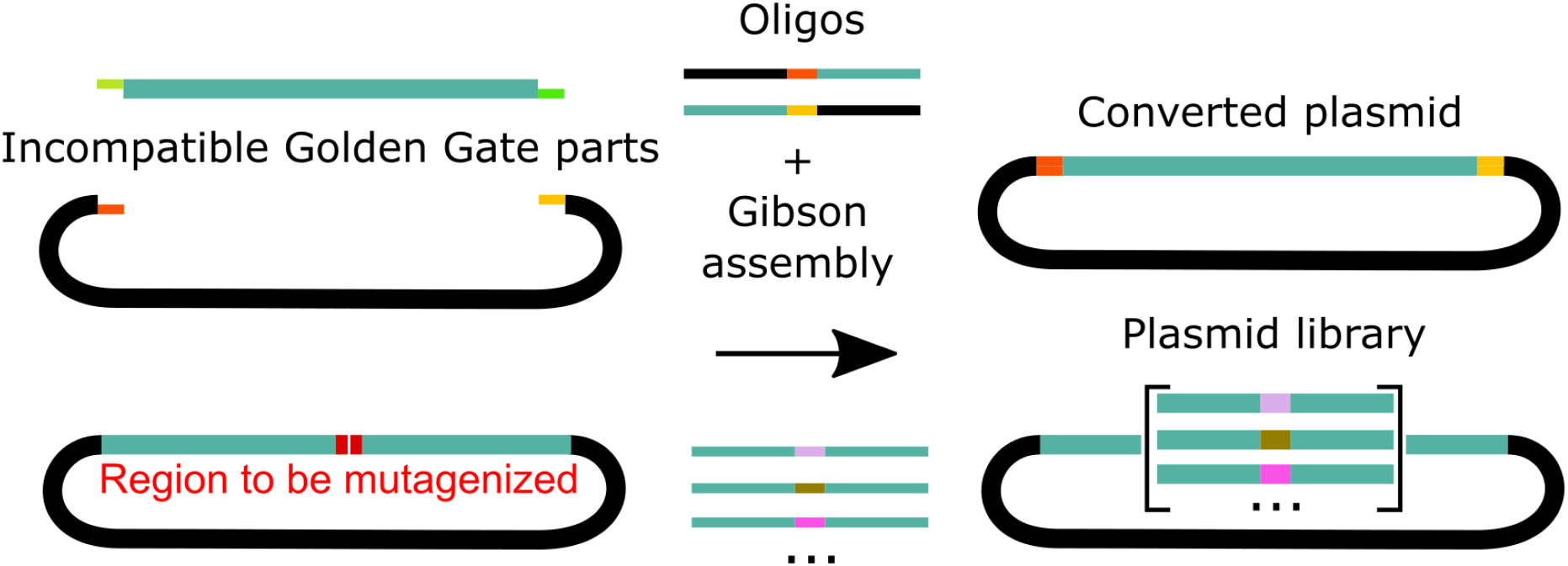

## Introduction

Molecular cloning is a cornerstone of modern biological research. One technique that has gained popularity in recent years is Gibson assembly^1^, which allows for seamless joining of multiple parts in a single reaction using three enzymes: 5’ exonuclease, DNA polymerase and DNA ligase. The only requirement is the presence of homologous regions (≥15 nucleotides) between parts. While Gibson assembly can be used to assemble transcriptional subunits following a common assembly standard^2,3^ the large cloning scars left between parts are often not desirable. Another drawback is the difficulty of correctly joining repetitive elements. Given these limitations, another technique reigns supreme where modular assembly is required, namely Golden Gate cloning. This technique is based on Type IIS restriction enzymes and a DNA ligase to join parts together. Because the enzymes typically leave 3-4 nt overhangs, only 1-2 amino acid scars are present in the final assemblies^4^. Many assembly standards have been described for use in prokaryotes and eukaryotes, which, unfortunately, are largely incompatible^5–9^. Assembly standards are usually swapped from one to another by PCR, but this requires multiple hands-on steps and may not be straightforward for large and/or complex parts.

While Gibson and Golden Gate methods are often depicted as being completely different molecular cloning paradigms, we find that there are many advantages to using both techniques in conjunction for modular assembly. In our lab for example, we typically use Gibson assembly for creating entry vectors. The price of the reaction is similar to that of Golden Gate cloning but can be completed in just 15 minutes. Furthermore, the complex toolsets of some of the standards^6,7^ can be reduced to just a single empty entry vector when using Gibson assembly: as the 5’ overhangs are digested by the exonuclease, sequences can be encoded in the insert PCR product to install the desired overhangs. Other reported - but underappreciated - properties of Gibson assembly are the ability to assemble DNA oligos^10,11^ (oligo stitching) and chew back of mismatched 3’ ends. Utilizing these features, we show that oligo stitching can readily convert Golden Gate entry vectors from one assembly standard into another with or without additional N/C-terminal fusions, directly assemble incompatible Golden Gate parts, generate *de novo* synthetic sequences, and be used for restriction enzyme site removal and plasmid saturation mutagenesis.

## Results and Discussion

### Clone conversion

We first set out to test the usefulness of using oligo stitching to convert one assembly standard into another (Figure 1A; Supplementary Table 1) by converting four vectors of the MoClo Toolkit^6^ into the GreenGate standard^7^. We used NEBuilder HiFi master mix with two 44-nt oligos, one for the left and one for the right flank, together with equal ratios of digested MoClo donor plasmid and GreenGate empty entry plasmid pGGC000 as the acceptor and sequenced five clones per assembly. The oligos match 20 bp of the backbone on one end and 20 bp of the part on the other end, with the 4 bases in the middle forming the new Golden Gate overhang (Figure 1A). We obtained hundreds to thousands of clones per assembly using homemade chemically-competent DH5α cells. The resulting cloning efficiency – i.e., the ratio of correct clones - ranged from 80 to 100%. The few clones containing errors all had a single deletion in the region specified by the oligo and all internal sequences were correct (Figure 1B; Supplementary Table 1). We also tested all possible orientations of the oligos and confirmed that all assemblies are equally efficient and accurate (Supplementary Table 1), indicating that the orientation of the oligos does not affect cloning efficiency, as could be expected based on the mechanism (Supplementary Figure 1).

**Figure 1.**
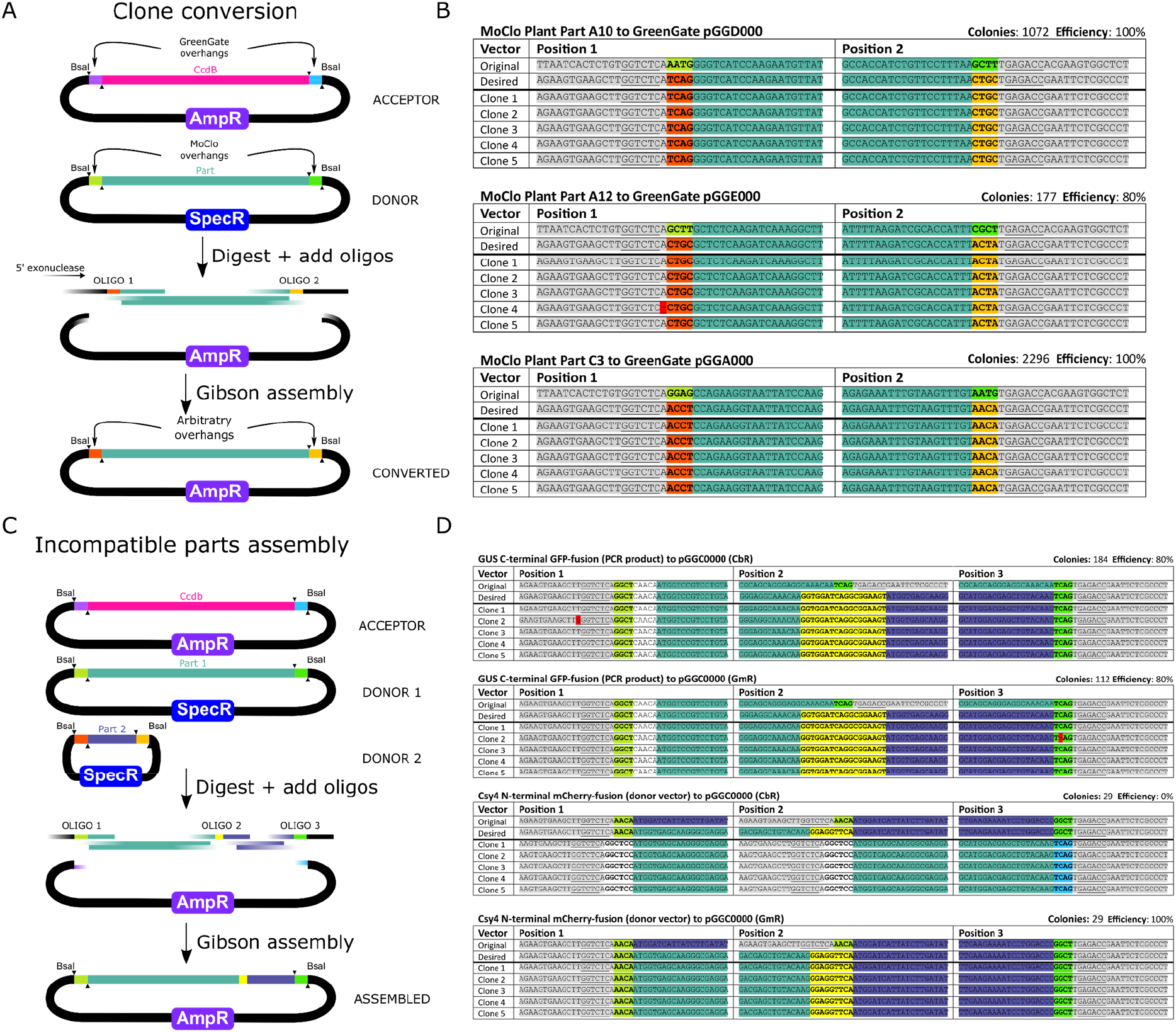
Clone conversion and incompatible parts assemblies. (A) In clone conversion GreenGate acceptor and MoClo donor plasmids are digested with BsaI and combined with two oligos containing 20-nt homologous sequences. Golden Gate overhangs are shown as colored sections directly adjacent to the black backbone and teal part. Via Gibson assembly the insert is transferred to the backbone of the acceptor plasmid with the new overhangs installed. Note that the overhangs of the final product (orange and yellow boxes) can be arbitrarily chosen. (B) Three representative clone conversion experiments converting MoClo parts into the GreenGate standard. The original and desired sequences are aligned at the top and the actual sequences of five randomly selected clones from each assembly. Each position shown corresponds to the stitching oligos used in the assembly. Mutations that deviate from the desired sequence are highlighted in red. (C) Incompatible parts assembly is an extension to the previous case, but with *n*+1 oligos (where *n* equals the number of parts). This can be used to combine incompatible Golden Gate parts derived from plasmids, PCR or oligos. (D) Two representative incompatible parts assemblies with the same carbenicillin-resistant (CbR) backbone as the donor or the gentamycin resistant (GmR) backbone. Each position shown corresponds to the stitching oligos used in the assembly. Yellow highlighted sequences are linkers encoded by oligo 2. Mutations deviating from the desired sequence are highlighted in red.

Another interesting case is converting overhangs *within* an assembly standard *(e.g.,* changing a C-terminal linker into a N-terminal linker). We selected four GreenGate vectors and again attempted to change the overhangs using two different oligos. However, we found carryover of the donor plasmid in the MtU6 assemblies, making it difficult to pick up desired clones (Supplementary Table 2). Similarly, when constructing part fusions (described below), we also found clones with the original donor plasmid in two out of three assemblies, with the worst case containing only the original donor plasmid (Supplementary Table 4). The original donor plasmid can be removed by gel extracting the insert, but this is not amenable to high-throughput conversion efforts. As an alternative, we constructed three GreenGate empty acceptor vectors containing gentamycin, tetracycline or spectinomycin resistance so that we can swap resistance markers and select against the contaminating donor plasmid. With the new acceptor vectors the overall efficiency increased; in the best case an increase of 0 to 100% correct colonies as compared to the previous experiment (Supplementary Table 4). For the MtU6 conversion experiments, there was essentially no difference in cloning efficiency between the original and the gentamycin backbone (Supplementary Table 2). However, the overhangs were actually correct in 80% of the clones, but the clones were not completely correct due to indels or substitutions.

To further explore the versatility of oligo stitching for clone conversion, we attempted to not only exchange overhangs, but also the type IIS restriction enzyme recognition sites. This type of modification relies on the 3’-5’ exonuclease proofreading activity of the DNA polymerase in the NEBuilder mix which can reportedly chew back up to 10-nt 3’ mismatches^12^ (Supplementary Figure 1B). To this end we changed the BsaI sites into AarI or SapI and moved these parts to the gentamycin resistance backbone, obtaining cloning efficiencies of 80-100% (Supplementary Table 3).

### Incompatible parts assembly

While it is advisable to convert all entry clones to the assembly standard one wants to use, in some cases it may be preferable to simply generate the expression vector directly from incompatible parts. PaperClip is one method that can be used for this, but it leaves an alanine scar at each junction^13^. To further test oligo stitching we selected a design where all overhangs are incompatible with each other: a MoClo promoter, a GreenGate coding sequence, terminator and 11-kb binary vector, for a total of three parts plus the backbone (Supplementary Figure 2A, B). All parts and the backbone were individually digested with BsaI and used for Gibson assembly with four stitching oligos. We only obtained eight colonies in homemade cells, but 100% of the five randomly selected clones were correct (Supplementary Table 4; Supplementary Figure 2C).

It is often desirable to mix and match parts in a single entry vector to perform an optimization experiment or to reduce the number of elements once an ideal architecture has been created *(e.g.* linkers, localization signals, *etc.).* Such parts are abundant in collections, but often contained in incompatible elements or lacking slight variations needed for the experiment. Therefore, we tested the ability to use oligo stitching to seamlessly and easily create three novel fusion parts using a combination of donor- or PCR-derived parts (Figure 1C). This approach was also highly successful; when using the gentamycin resistance backbone, we obtained 80-100% correct clones (Figure 1D, Supplementary Table 4).

In some cases it is useful to have a given part at all or many of the possible positions in a Golden Gate assembly scheme (*e.g.*, linkers, fluorescent proteins, tags, *etc.).* When these parts are not yet present in a Golden Gate format or lack adjacent restriction enzyme recognition sites, PCR followed by cleanup or gel extraction is needed to generate the parts. To avoid amplification and cleanup of each separate PCR product for each position, we reasoned that it might be more efficient to generate one single PCR product *without* overhangs and then assemble this amplicon into an empty entry vector with two oligos specifying the overhangs. To test this, we amplified the glutathione-*S*-transferase (GST) and 3 x hemagglutinin (3xHA) tags from two different Gateway vectors with a single primer pair for each amplicon. We then assembled these amplicons into five GreenGate positions using five sets of DNA oligos and obtained a cloning efficiencies of 80-100% in nine out of ten cases, with the remaining case having an efficiency of 60% (Supplementary Table 5).

We also tested the ability to add relatively small sequences via oligos as opposed to PCR products or donor vectors. An ER-peptide signal (66 bp) and a SV40 nuclear localization signal (21 bp) were added to the left and right oligos, respectively, to fuse this to one of the parts. The three possible combinations (i.e., N/C/NC-terminal fusions) showed cloning efficiencies of 80-100% (Supplementary Table 6).

### De novo part assembly

The ability of Gibson assembly to convert DNA oligos into synthetic dsDNA sequences was recognized early^14^, and has been used for, among others, the construction of prime editing guide RNAs (pegRNAs) for prime editing^15^. Here we expand on those results by showing that we can increase the number of oligos in the reaction to additionally tag the pegRNA with other elements such as the 5’ *FLOWERING LOCUS T* (*FT*) cell-to-cell mobile signal^16^ and/or a 3’ structured RNA motif to resist degradation^17^ by making use of four or five oligos. The cloning yield dropped with five oligos, but the cloning efficiency was 60-100% for all four designs (Figure 2B), making this a viable approach to building DNA fragments at least ~270 bp long.

**Figure 2.**
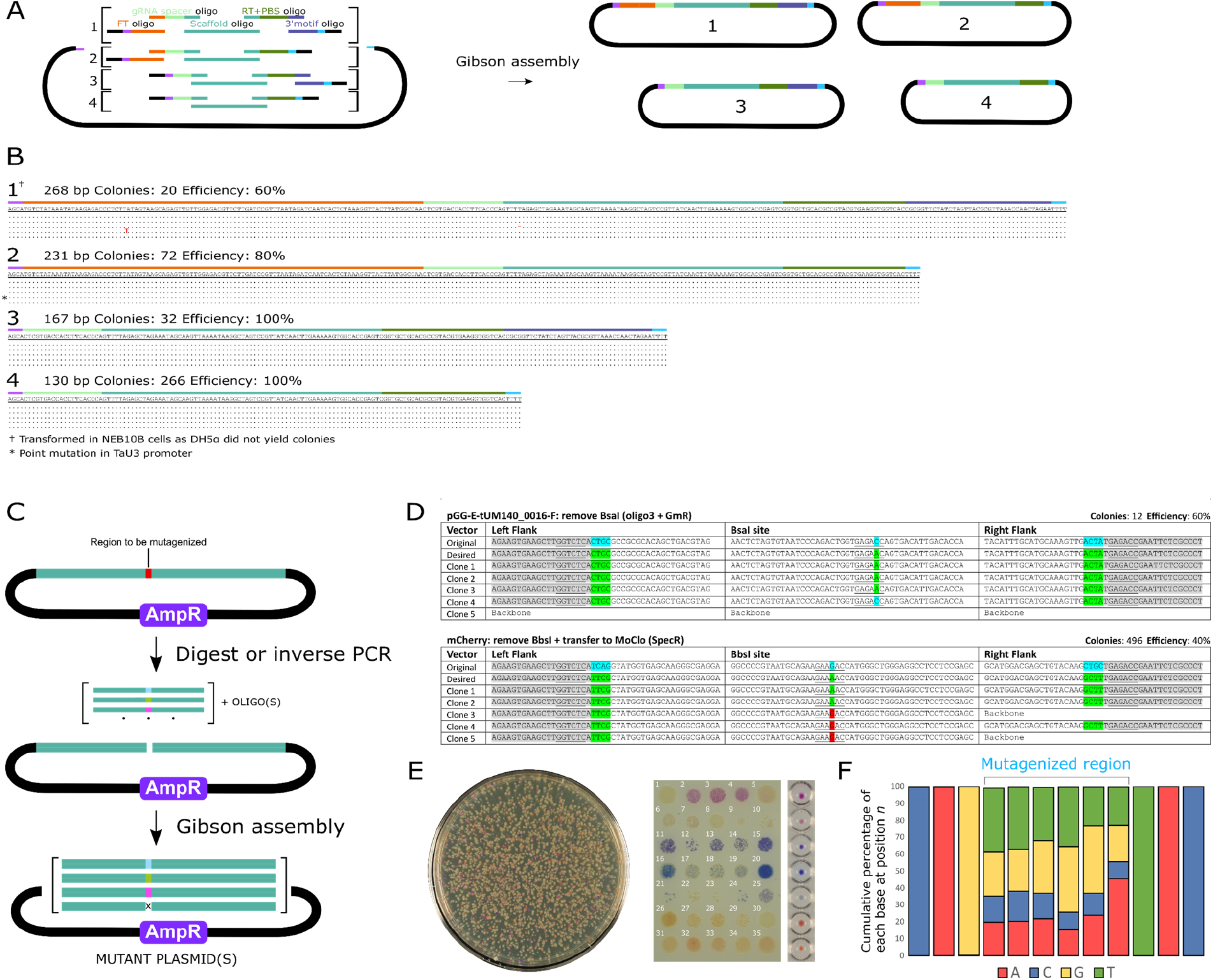
*De novo* synthetic sequences and restriction enzyme mutagenesis. (A) Three to five oligos are mixed with digested backbone and assembly mix to generate one single pegRNA per reaction. The orientation of the oligos alternates between sense and antisense as previously described^14^. (B) Alignments of the expected and observed sequences for five randomly picked clones for each of the four assemblies. Dots are used to indicate identical bases, deviating bases are shown in red. (C) Restriction enzyme recognition sequence removal and saturation mutagenesis. The red position indicates the region targeted for mutagenesis. The sequence is linearized at this position either by restriction digest or by inverse PCR and then mixed with an oligo for site-directed mutagenesis or with a pool of degenerate oligos for site saturation mutagenesis. (D) Representative examples of clones domesticated for BsaI and BbsI. Each position shown corresponds to the stitching oligos used in the assembly. Mutations that deviate from the desired sequence are highlighted in red. (E) Left: Representative Petri dish after transformation in chemically competent *E. coli* cells with amilCP_Orange mutagenesis assembly. Right: Colonies selected by color and genotyped as shown in Supplementary Table 8. Each row contains five independent clones of the same color from the transformation plate. The right column shows the representative colors from pelleted cells. (F) Cumulative percentage of bases for each position at the mutagenized region ±3 bases.

### Restriction enzyme removal and saturation mutagenesis

When swapping DNA parts between assembly standards, it is often also necessary to remove internal restriction enzyme sites in a process called domestication. Therefore we tested oligo stitching to domesticate vectors by removing either XbaI or EcoRI, two Type II enzymes used in BioBrick assembly^18^, again relying on the proofreading activity of the DNA polymerase. The vectors were digested with their respective enzymes and assembled with DNA oligos encoding silent mutations. As opposed to the previous N/C terminal fusion results with BsaI digestion, our five randomly selected clones were all recombinant plasmids, suggesting that the efficiency of the restriction enzyme may play a role in preventing the carryover of donor plasmid. Cloning efficiencies ranged between 60 and 100%, again with errors only being present in the region specified by the oligo. We then attempted Type IIS recognition site removal (BsaI, BbsI and AarI) by digesting the plasmids with their respective enzymes and assembling with oligos encoding silent mutations and the Golden Gate flanking sequences (Figure 2D). Success was variable, with cloning efficiencies between 0 and 60% (Supplementary Table 7). By altering either the backbone or the position of the silent mutation in the mutagenic oligo, we could domesticate all parts with a minimum cloning efficiency of 20%, indicating that domestication of Type IIS sites is feasible, but more clones might be needed (Supplementary Table 7).

Motivated by these results, we also tested if plasmid saturation mutagenesis is a feasible application. To enable a visual readout of mutagenesis, we made use of an amilCP_Orange chromoprotein encoding vector as mutagenesis of just six nucleotides (two amino acids) can alter the orange color of the chromoprotein^19^. Conveniently, the restriction enzyme PflMI cuts close to the position to be mutagenized, requiring a 7-nt 3’-end chew-back to remove the codons encoding these two amino acids (Supplementary Figure 3A). After gel extraction of the digested vector, we combined a mutagenic oligo containing the degenerate “NNNNNN” identities flanked by 20-bp of sequence matching the amilCP_Orange CDS on both sides (Supplementary Figure 3A). We recovered approximately 10,000 colonies with seven distinct colors we could discern by eye. The majority of the corresponding amino acid sequences were not reported in the mutagenesis screen of the original publication (Supplementary Table 8). Upon restreaking, there was a high occurrence of the loss of chromoprotein expression (Supplementary Figure 2E), in line with previous observations^19^. We performed amplicon sequencing of a pool of approximately 80,000 colonies to better characterize the mutagenesis screen. The original vector sequence was present in less than 3.4% of the reads and all other reads showed variable nucleotides at the position specified by the mutagenic oligo. The nucleotide identities at these positions were slightly biased against cytosine (Supplementary Figure 3F). Bias in oligonucleotide synthesis is a well-known phenomenon^20^, but biological selection may also have contributed to this observation. Deletions were observed in only 0.72% of the reads in the region specified by the mutagenic oligo. For all other reads the correct nucleotide at each position was detected in at least 99.7% of the reads. At the DNA level, 99.5% of all possible 4,096 hexanucleotide variants were present in the pool. Interestingly, variants encoding stop codons were enriched for, which is in line with the observation that chromoprotein expression is selected against and reduces the growth rate^19^. At the protein level, all 400 possible amino acid combinations are represented, and expectedly, variants encoding at least one stop codon are also enriched (Supplementary Figure 3B,C).

In conclusion, we show that oligo stitching is a powerful, efficient and flexible tool to easily convert between cloning design standards and adapt or build DNA parts for Golden Gate assembly. One limitation of our approach is the reliance on restriction enzyme recognition sites to linearize the DNA acceptors and donors. However, the combination of CRISPR/Cas cloning reagents and oligo stitching could open up an effectively unlimited design space within plasmids. Ultimately, we anticipate that oligo stitching could be used to convert entire part collections *en masse* allowing for more widespread sharing and reuse of parts regardless of the assembly standard.

## Methods

### Plasmids

The MoClo Toolkit (Addgene Kit #1000000044), MoClo Plant Parts Kit (Addgene Kit #1000000047), GreenGate Cloning System (Addgene Kit #1000000036) and amilCP_Orange chromoprotein vector (Addgene Plasmid #117850) were acquired from Addgene. The plasmids pEN-R2-GST-L3 and pEN-R2-3xHA-L3 were previously published^22^. The novel plasmids pGGC000-GmR, pGGC000-SpecR and pGGC000-TetR created here are available for distribution via https://gatewayvectors.vib.be/.

### Plasmid preparation

All plasmids were extracted using the GeneJET Plasmid MiniPrep Kit (Thermo Fisher) according to the manufacturer’s instructions. For the assembly standard conversion experiments, all MoClo and GreenGate vectors were individually digested with 0.5 μL BsaI-HF®v2 (NEB) with CutSmart buffer (NEB) in a reaction volume of 10 μL for 4 hours at 37°C, followed by an inactivation step at 80°C for 20 min. Digested vectors were stored at −20°C.

For the removal of restriction enzyme recognition sites of XbaI and EcoRI, the vectors were digested with 0.5 μL of the respective Promega enzyme, 2 μL BSA (1 mg/mL) and 2 μL buffer H (Promega) in reaction volume of 20 μL. The other conditions were the same as described above.

For saturation mutagenesis experiments, 1 μg of amilCP_Orange chromoprotein was digested with 1 μL PflMI (NEB) with r3.1 buffer (NEB) in a reaction volume of 50 μL for 4 hours at 37°C, followed by an inactivation step at 80°C for 20 min. The reaction was run on an 0.8% agarose gel stained with SybrSafe and the band corresponding to the linearized vector was excised under blue safety light. Gel extraction was done using the Zymoclean Gel DNA Recovery Kit (Zymo Research) according to the manufacturer’s instructions.

### PCR amplicon preparation

For experiment 32, GFP was amplified using primers GFP_StitAmp_F and GFP_StitAmp_R with Q5 polymerase from pGG-D-GFP-E (pGGD001) using the following PCR conditions: 98°C/5 min + 30 x (98°C/10 sec + 60°C/30 sec + 72°C/30 sec) + 72°C/5 min + 23°C/∞. The fragment of the correct size was purified using the Zymoclean Gel DNA Recovery Kit. For experiments 13-17 and 18-22, pEN-R2-GST-L3 and pEN-R2-3xHA-L3 was used as the template with primers 3xHA_FW/REV and GST_FW/REV, respectively. The same PCR conditions were used as for GFP amplification. No PCR cleanup was done for these two samples.

### NEBuilder assembly

Each assembly reaction was run in a reaction volume of 10 μL, with half of the volume composed of NEBuilder master mix (NEB) according to a previously reported protocol with modifications^11^. 0.04 pmol was used for each of the insert(s) and the backbone. Oligos were designed to have homology with 20 bp at each side of the junction. Extra sequences such as overhangs, stop codons or linkers were included when necessary. Oligos were resuspended to 100 μM and diluted to 0.3 pmol/μl (see Supplementary protocol). The reaction was incubated at 50°C for 1h and put on ice until *E. coli* transformation. An overview of the oligos can be found in Supplementary Table 9.

For the one PCR to many entries experiments, 1 μL of unpurified PCR product, 1 uL of digested pGGC000, 1 μL of oligo mix, 2 μL of water and 5 μL of NEBuilder was used. The rest of the conditions were the same as described above. For the assembly of incompatible library parts, we made use of 1 μL of each digested part and backbone vector with a concentration of 50 ng/μL. The rest of the conditions were the same as described above.

For the synthesis of pegRNAs we made use of a previously described protocol^15^. Briefly, 50 ng of digested pGG-TaU3-ccdB-T(n7) was assembled with 1 μL of each 100 nM oligo in a 10 μL NEBuilder assembly reaction.

### *E. coli* transformation

We made use of home-made DH5α chemically competent cells with a measured transformation efficiency of 4.5×10^6^ cfu/μg. 2 μL of the assembly mix was mixed with 25 μL of competent cells in an ice-cold 1.5 mL Eppendorf tube. After 30 min incubation on ice, the reaction was heat shocked for 30 seconds at 42°C and chilled on ice for 5 min. 300 μL SOC was added, and the tube was incubated at 37°C for 60 minutes in a shaking incubator. 100 μL was plated on pre-warmed (37°C) LB medium containing the appropriate antibiotics.

For electrocompetent cells we made use of NEB 10β with a transformation efficiency of 2×10^10^ cfu/μg. One microliter of the assembly reaction was mixed with 50 μL of competent cells and placed inside a chilled electroporation cuvette (0.2 cm gap, BioRad). The electroporation was carried out in a GenePulser (BioRad) according to the manufacturer’s conditions and 900 μL of SOC was added immediately to the cells. Cells were incubated at 37°C for 60 minutes in a shaking incubator. 100 μL was plated on pre-warmed (37°C) LB medium containing the appropriate antibiotics.

### Analysis of clones

Single *E. coli* colonies were picked and grown overnight in 3 mL of LB medium containing the appropiate antibiotics. Plasmids were extracted and sent for Sanger sequencing (Mix2Seq, Eurofins, Germany) using the appropriate primers (Supplementary Table 9).

### Sanger and NGS analysis of saturation mutagenesis experiment

Several clones of each color were transferred to new plates for archiving. A toothpick was used to pick up bacterial material of each clone which was then lysed in 15 μL lysis buffer (10 mM Tris and 0.1% Triton X-100) by boiling the reaction for 5 min. The reaction was then spun down for 5 min at 4000 rpm. For the PCR reaction, 2 μL of the supernatant was used in a REDTaq ReadyMix PCR Reaction with primers Orange_MUT_seq_F and R (Supplementary Table 9). The PCR reactions were cleaned up with HighPrep™ PCR Clean-up beads (MagBio) according to the manufacturer’s instructions and sent for Sanger sequencing (Eurofins, Germany) with primer Orange_MUT_seq_F.

For the NGS analysis, we collected all colonies from the NEB 10β transformed cells by carefully scraping the LB medium in the presence of 5 mL LB medium. The medium was pooled of all these plates and divided into two. Each tube was processed with the ZymoPURE II Plasmid Midiprep Kit (Zymo Research). The resulting midipreps were pooled and a dilution of 10 ng/μL was prepared. Eight separate 50 μL PCR reactions were set up with Q5 (NEB) using 2.5 μL of the diluted midiprep as the template and with the primers Orange_NGS_F and R. The PCR conditions used were as follows: 98°C/5 min + 12 x (98°C/10 sec + 68°C/30 sec + 72°C/20 sec) + 72°C/5 min + 23°C/∞. The PCR reactions were pooled and purified using the GeneJET PCR Purification Kit (Thermo Fisher) according to the manufacturer’s instructions. The sample was then sent to Eurofins (Germany) for adapter ligation and NGS sequencing (5 million paired reads, 2×150 bp). The data was analyzed using CRISPResso2^23^. The guide sequence was set as “ACCACAGGTTGGATACGGAA”, the minimum average read quality to a Phred score of 30, and the window size as 26 with the quantification center in the middle of the degenerate hexanucleotide. Allelic variants with less than 10 reads were filtered out before further data analysis in Excel.

## Supporting information

Supplementary protocol

## Supporting information

Additional information references in the text are tables showing the sequences of the recombinant clones (Supplementary Tables 1-8) and a table with the oligo sequences (Supplementary Table 9). Supplementary Figure 1 shows the mechanism of the reaction. Supplementary Figure 2 shows more information concerning the incompatible library experiment. Supplementary Figure 3 shows an overview of the saturation mutagenesis experiment. The supplementary protocol is designed to provide a step-by-step protocol to be used in the lab.

## Author contributions

J.D.S., M.V. and T.B.J. conceptualized and designed the research; J.D.S. and M.V. performed the wet lab experiments. J.D.S. and C.G. performed the NGS analysis. J.D.S., and T.B.J. drafted the manuscript, J.D.S., M.V., C.G. and T.B.J. critically reviewed the manuscript. All authors read and approved the final manuscript.

## Acknowledgements

We thank Sylvestre Marillonnet for sharing the MoClo Toolkit (Addgene Kit #1000000044), Nicola Patron for sharing the MoClo Plant Parts Kit (Addgene Kit #1000000047) and Anthony Forster for sharing the amilCP_Orange chromoprotein vector (Addgene Plasmid #117850). We thank Jan Lohmann for sharing the GreenGate Cloning System (Addgene Kit #1000000036). Karel Spruyt is thanked for taking the photographs of the bacterial plates. J.D.S is supported by Ghent University (‘Bijzonder Onderzoeksfonds Methusalem project’ no. BOF15/MET_V/004). C.G. is supported by VLAIO (Flanders Innovation & Entrepreneurship) WHEAT-BEG (project number HBC.2018.2152)

## Notes

The authors declare no competing interests.

## Notes

### Competing Interest Statement

The authors have declared no competing interest.

## References

1. Gibson, D. G. et al. Enzymatic assembly of DNA molecules up to several hundred kilobases. Nat. Methods 6, 343–345 (2009).

2. Akama-Garren, E. H. et al. A Modular Assembly Platform for Rapid Generation of DNA Constructs. Sci. Rep. 6, 16836 (2016).

3. Torella, J. P. et al. Unique nucleotide sequence-guided assembly of repetitive DNA parts for synthetic biology applications. Nat. Protoc. 9, 2075–2089 (2014).

4. Marillonnet, S. & Grützner, R. Synthetic DNA Assembly Using Golden Gate Cloning and the Hierarchical Modular Cloning Pipeline. Curr. Protoc. Mol. Biol. 130, e115 (2020).

5. Moore, S. J. et al. EcoFlex: A Multifunctional MoClo Kit for E. coli Synthetic Biology. ACS Synth. Biol. 5, 1059–1069 (2016).

6. Weber, E., Engler, C., Gruetzner, R., Werner, S. & Marillonnet, S. A Modular Cloning System for Standardized Assembly of Multigene Constructs. PLoS One 6, e16765 (2011).

7. Lampropoulos, A. et al. GreenGate - A Novel, Versatile, and Efficient Cloning System for Plant Transgenesis. PLoS One 8, e83043 (2013).

8. Chiasson, D. et al. A unified multi-kingdom Golden Gate cloning platform. Sci. Rep. 9, 10131 (2019).

9. Patron, N. J. et al. Standards for plant synthetic biology: a common syntax for exchange of DNA parts. New Phytol. 208, 13–19 (2015).

10. Jacobs, T. B. & Martin, G. B. High-throughput CRISPR Vector Construction and Characterization of DNA Modifications by Generation of Tomato Hairy Roots. JoVE e53843 (2016) doi:doi:10.3791/53843.

11. Utilizing both homology and oligonucleotide stitching techniques to build large constructs. ThermoFisher Scientific Inc. white paper. https://assets.thermofisher.com/TFS-Assets/BID/Reference-Materials/gibson-assembly-hifi-homology-oligonucleotide-stitching-white-paper.pdf (2020). Accessed February 2022.

12. Hsieh, P. Bridging dsDNA with a ssDNA Oligo and NEBuilder® HiFi DNA Assembly to create an sgRNA-Cas9 Expression Vector. New England Biolabs Inc. Application Note. https://www.neb.com/-/media/nebus/files/application-notes/appnote_bridging_dsdna_with_ssdna_oligo_and_nebuilder_hifi_dna_assembly_to_create_sgrna-cas9_expression_vector.pdf. (2018) Accessed February 2022.

13. Trubitsyna, M., Michlewski, G., Cai, Y., Elfick, A. & French, C. E. PaperClip: rapid multi-part DNA assembly from existing libraries. Nucleic Acids Res. 42, e154–e154 (2014).

14. Gibson, D. G., Smith, H. O., Hutchison, C. A., Venter, J. C. & Merryman, C. Chemical synthesis of the mouse mitochondrial genome. Nat. Methods 7, 901–903 (2010).

15. Kweon, J. et al. Engineered prime editors with PAM flexibility. Mol. Ther. 29, 2001–2007 (2021).

16. Lei, J. et al. Heritable gene editing using FT mobile guide RNAs and DNA viruses. Plant Methods 17, 20 (2021).

17. Nelson, J. W. et al. Engineered pegRNAs improve prime editing efficiency. Nat. Biotechnol. (2021) doi:10.1038/s41587-021-01039-7.

18. Knight, T. F. Idempotent Vector Design for Standard Assembly of BioBricks. Tech. rep., MIT Synth. Biol. Work. Gr. Tech. Reports (2003).

19. Liljeruhm, J. et al. Engineering a palette of eukaryotic chromoproteins for bacterial synthetic biology. J. Biol. Eng. 12, 8 (2018).

20. Kawai, F., Nakamura, A., Visootsat, A. & Iino, R. Plasmid-Based One-Pot Saturation Mutagenesis and Robot-Based Automated Screening for Protein Engineering. ACS Omega 3, 7715–7726 (2018).

21. Wang, J.W. et al. CRISPR/Cas9 nuclease cleavage combined with Gibson assembly for seamless cloning. Biotechniques 58, 161–170 (2015).

22. Karimi, M., De Meyer, B. & Hilson, P. Modular cloning in plant cells. Trends Plant Sci. 10, 103–105 (2005).

23. Clement, K. et al. CRISPResso2 provides accurate and rapid genome editing sequence analysis. Nat. Biotechnol. 37, 224–226 (2019).

