## Supplementary Table for "Simple and efficient modification of Golden Gate design standards and parts using oligo stitching"

**SUPPLEMENTARY MATERIALS**

### Supplementary Table 1: Clone conversion

Partial sequences of the original sequence, the desired sequence and five sequenced clones are shown in each table. The BsaI recognition motif is underlined. The 4-bp overhangs that are generated by BsaI digestion are shown in bold, flanked by 20 bp of sequence at both sides. The overhang is shown in blue for the original MoClo sequence and in green for the desired GreenGate sequence. Additional bases that were added, such as start codons or bases needed to keep the sequences in frame after assembly are also highlighted in green. The sequence highlighted in gray is the backbone sequence. Mutations that deviate from the desired sequence are highlighted in red. The number of colonies and the cloning efficiency is also indicated. Experiment number is indicated in top left corner.

**1) MoClo Plant Part A10 (CDS1 module) to GreenGate pGGD000**
**Colonies**: 1072 / **Efficiency**: 100%

| Vector | Position 1 | Position 2 |
|---|---|---|
| Original | TTAATCACTCTGTGGTCTCA**AATG**GGGTCATCCAAGAATGTTAT | GCCACCATCTGTTCCTTTAA**GCTT**TGAGACCACGAAGTGGCTCT |
| Desired | AGAAGTGAAGCTTGGTCTCA**TCAG**GGGTCATCCAAGAATGTTAT | GCCACCATCTGTTCCTTTAA**CTGC**TGAGACCGAATTCTCGCCCT |
| Clone 1 | AGAAGTGAAGCTTGGTCTCA**TCAG**GGGTCATCCAAGAATGTTAT | GCCACCATCTGTTCCTTTAA**CTGC**TGAGACCGAATTCTCGCCCT |
| Clone 2 | AGAAGTGAAGCTTGGTCTCA**TCAG**GGGTCATCCAAGAATGTTAT | GCCACCATCTGTTCCTTTAA**CTGC**TGAGACCGAATTCTCGCCCT |
| Clone 3 | AGAAGTGAAGCTTGGTCTCA**TCAG**GGGTCATCCAAGAATGTTAT | GCCACCATCTGTTCCTTTAA**CTGC**TGAGACCGAATTCTCGCCCT |
| Clone 4 | AGAAGTGAAGCTTGGTCTCA**TCAG**GGGTCATCCAAGAATGTTAT | GCCACCATCTGTTCCTTTAA**CTGC**TGAGACCGAATTCTCGCCCT |
| Clone 5 | AGAAGTGAAGCTTGGTCTCA**TCAG**GGGTCATCCAAGAATGTTAT | GCCACCATCTGTTCCTTTAA**CTGC**TGAGACCGAATTCTCGCCCT |

**2) MoClo Plant Part A12 (3U + Ter module) to GreenGate pGGE000**
**Colonies**: 177 / **Efficiency**: 80%

| Vector | Position 1 | Position 2 |
|---|---|---|
| Original | TTAATCACTCTGTGGTCTCA**GCTT**GCTCTCAAGATCAAAGGCTT | ATTTTAAGATCGCACCATTT**CGCT**TGAGACCACGAAGTGGCTCT |
| Desired | AGAAGTGAAGCTTGGTCTCA**CTGC**GCTCTCAAGATCAAAGGCTT | ATTTTAAGATCGCACCATTT**ACTA**TGAGACCGAATTCTCGCCCT |
| Clone 1 | AGAAGTGAAGCTTGGTCTCA**CTGC**GCTCTCAAGATCAAAGGCTT | ATTTTAAGATCGCACCATTT**ACTA**TGAGACCGAATTCTCGCCCT |
| Clone 2 | AGAAGTGAAGCTTGGTCTCA**CTGC**GCTCTCAAGATCAAAGGCTT | ATTTTAAGATCGCACCATTT**ACTA**TGAGACCGAATTCTCGCCCT |
| Clone 3 | AGAAGTGAAGCTTGGTCTCA**CTGC**GCTCTCAAGATCAAAGGCTT | ATTTTAAGATCGCACCATTT**ACTA**TGAGACCGAATTCTCGCCCT |
| Clone 4 | AGAAGTGAAGCTTGGTCTC-**CTGC**GCTCTCAAGATCAAAGGCTT | ATTTTAAGATCGCACCATTT**ACTA**TGAGACCGAATTCTCGCCCT |
| Clone 5 | AGAAGTGAAGCTTGGTCTCA**CTGC**GCTCTCAAGATCAAAGGCTT | ATTTTAAGATCGCACCATTT**ACTA**TGAGACCGAATTCTCGCCCT |

**3) MoClo Plant Part C3 (PRO + 5U module) to GreenGate pGGA000**
**Colonies**: 2296 / **Efficiency**: 100%

| Vector | Position 1 | Position 2 |
|---|---|---|
| Original | TTAATCACTCTGTGGTCTCA**GGAG**CCAGAAGGTAATTATCCAAG | AGAGAAATTTGTAAGTTTGT**AATG**TGAGACCACGAAGTGGCTCT |
| Desired | AGAAGTGAAGCTTGGTCTCA**ACCT**CCAGAAGGTAATTATCCAAG | AGAGAAATTTGTAAGTTTGT**AACA**TGAGACCGAATTCTCGCCCT |
| Clone 1 | AGAAGTGAAGCTTGGTCTCA**ACCT**CCAGAAGGTAATTATCCAAG | AGAGAAATTTGTAAGTTTGT**AACA**TGAGACCGAATTCTCGCCCT |
| Clone 2 | AGAAGTGAAGCTTGGTCTCA**ACCT**CCAGAAGGTAATTATCCAAG | AGAGAAATTTGTAAGTTTGT**AACA**TGAGACCGAATTCTCGCCCT |
| Clone 3 | AGAAGTGAAGCTTGGTCTCA**ACCT**CCAGAAGGTAATTATCCAAG | AGAGAAATTTGTAAGTTTGT**AACA**TGAGACCGAATTCTCGCCCT |

### SUPPLEMENTARY MATERIALS

| | | |
|---|---|---|
| Clone 4 | AGAAGTGAAGCTTGGTCTCA**ACCT**CCAGAAGGTAATTATCCAAG | AGAGAAATTTGTAAGTTTGT**AACA**TGAGACCGAATTCTCGCCCT |
| Clone 5 | AGAAGTGAAGCTTGGTCTCA**ACCT**CCAGAAGGTAATTATCCAAG | AGAGAAATTTGTAAGTTTGT**AACA**TGAGACCGAATTCTCGCCCT |

**4) MoClo Plant Part C7 (NT1 module) to GreenGate pGGC000**

**Colonies**: 1476 / **Efficiency**: 80%

| Vector | Position 1 | Position 2 |
|---|---|---|
| Original | GAAGAGCCACTGTGGTCTCA**CCAT**GGTGAGCAAGGGCGAGGAGC | AAAAACGCGGCTATTAGATC**AATG**TGAGACCACGAGTGATTAAT |
| Desired | AGAAGTGAAGCTTGGTCTCA**GGCT**CCATGGTGAGCAAGGGCGAG | AAAACGCGGCTATTAGATCA**TCAG**TGAGACCGAATTCTCGCCCT |
| Clone 1 | AGAAGTGAAGCTTGGTCTCA**GGCT**CCATGGTGA-CAAGGGCGAG | AAAACGCGGCTATTAGATCA**TCAG**TGAGACCGAATTCTCGCCCT |
| Clone 2 | AGAAGTGAAGCTTGGTCTCA**GGCT**CCATGGTGAGCAAGGGCGAG | AAAACGCGGCTATTAGATCA**TCAG**TGAGACCGAATTCTCGCCCT |
| Clone 3 | AGAAGTGAAGCTTGGTCTCA**GGCT**CCATGGTGAGCAAGGGCGAG | AAAACGCGGCTATTAGATCA**TCAG**TGAGACCGAATTCTCGCCCT |
| Clone 4 | AGAAGTGAAGCTTGGTCTCA**GGCT**CCATGGTGAGCAAGGGCGAG | AAAACGCGGCTATTAGATCA**TCAG**TGAGACCGAATTCTCGCCCT |
| Clone 5 | AGAAGTGAAGCTTGGTCTCA**GGCT**CCATGGTGAGCAAGGGCGAG | AAAACGCGGCTATTAGATCA**TCAG**TGAGACCGAATTCTCGCCCT |

**5) MoClo Plant Part A10 (CDS1 module) to GreenGate pGGD000 (Sense/Sense)**

**Colonies:** 180 / **Efficiency**: 100%

| Vector | Position 1 | Position 2 |
|---|---|---|
| Original | TTAATCACTCTGTGGTCTCA**AATG**GGGTCATCCAAGAATGTTAT | GCCACCATCTGTTCCTTTAA**GCTT**TGAGACCACGAAGTGGCTCT |
| Desired | AGAAGTGAAGCTTGGTCTCA**TCAG**GGGTCATCCAAGAATGTTAT | GCCACCATCTGTTCCTTTAA**CTGC**TGAGACCGAATTCTCGCCCT |
| Clone 1 | AGAAGTGAAGCTTGGTCTCA**TCAG**GGGTCATCCAAGAATGTTAT | GCCACCATCTGTTCCTTTAA**CTGC**TGAGACCGAATTCTCGCCCT |
| Clone 2 | AGAAGTGAAGCTTGGTCTCA**TCAG**GGGTCATCCAAGAATGTTAT | GCCACCATCTGTTCCTTTAA**CTGC**TGAGACCGAATTCTCGCCCT |
| Clone 3 | AGAAGTGAAGCTTGGTCTCA**TCAG**GGGTCATCCAAGAATGTTAT | GCCACCATCTGTTCCTTTAA**CTGC**TGAGACCGAATTCTCGCCCT |
| Clone 4 | AGAAGTGAAGCTTGGTCTCA**TCAG**GGGTCATCCAAGAATGTTAT | GCCACCATCTGTTCCTTTAA**CTGC**TGAGACCGAATTCTCGCCCT |
| Clone 5 | AGAAGTGAAGCTTGGTCTCA**TCAG**GGGTCATCCAAGAATGTTAT | GCCACCATCTGTTCCTTTAA**CTGC**TGAGACCGAATTCTCGCCCT |

**6) MoClo Plant Part A10 (CDS1 module) to GreenGate pGGD000 (Antisense/Antisense)**

**Colonies:** 414 / **Efficiency**: 100%

| Vector | Position 1 | Position 2 |
|---|---|---|
| Original | TTAATCACTCTGTGGTCTCA**AATG**GGGTCATCCAAGAATGTTAT | GCCACCATCTGTTCCTTTAA**GCTT**TGAGACCACGAAGTGGCTCT |
| Desired | AGAAGTGAAGCTTGGTCTCA**TCAG**GGGTCATCCAAGAATGTTAT | GCCACCATCTGTTCCTTTAA**CTGC**TGAGACCGAATTCTCGCCCT |
| Clone 1 | AGAAGTGAAGCTTGGTCTCA**TCAG**GGGTCATCCAAGAATGTTAT | GCCACCATCTGTTCCTTTAA**CTGC**TGAGACCGAATTCTCGCCCT |
| Clone 2 | AGAAGTGAAGCTTGGTCTCA**TCAG**GGGTCATCCAAGAATGTTAT | GCCACCATCTGTTCCTTTAA**CTGC**TGAGACCGAATTCTCGCCCT |
| Clone 3 | AGAAGTGAAGCTTGGTCTCA**TCAG**GGGTCATCCAAGAATGTTAT | GCCACCATCTGTTCCTTTAA**CTGC**TGAGACCGAATTCTCGCCCT |
| Clone 4 | AGAAGTGAAGCTTGGTCTCA**TCAG**GGGTCATCCAAGAATGTTAT | GCCACCATCTGTTCCTTTAA**CTGC**TGAGACCGAATTCTCGCCCT |
| Clone 5 | AGAAGTGAAGCTTGGTCTCA**TCAG**GGGTCATCCAAGAATGTTAT | GCCACCATCTGTTCCTTTAA**CTGC**TGAGACCGAATTCTCGCCCT |

**7) MoClo Plant Part A10 (CDS1 module) to GreenGate pGGD000 (Sense/Antisense)**

**Colonies:** 136 / **Efficiency**: 100%

| Vector | Position 1 | Position 2 |
|---|---|---|
| Original | TTAATCACTCTGTGGTCTCA**AATG**GGGTCATCCAAGAATGTTAT | GCCACCATCTGTTCCTTTAA**GCTT**TGAGACCACGAAGTGGCTCT |
| Desired | AGAAGTGAAGCTTGGTCTCA**TCAG**GGGTCATCCAAGAATGTTAT | GCCACCATCTGTTCCTTTAA**CTGC**TGAGACCGAATTCTCGCCCT |
| Clone 1 | AGAAGTGAAGCTTGGTCTCA**TCAG**GGGTCATCCAAGAATGTTAT | GCCACCATCTGTTCCTTTAA**CTGC**TGAGACCGAATTCTCGCCCT |

| | | |
|---|---|---|
| Clone 2 | AGAAGTGAAGCTTGGTCTCA**TCAG**GGGTCATCCAAGAATGTTAT | GCCACCATCTGTTCCTTTAA**CTGC**TGAGACCGAATTCTCGCCCT |
| Clone 3 | AGAAGTGAAGCTTGGTCTCA**TCAG**GGGTCATCCAAGAATGTTAT | GCCACCATCTGTTCCTTTAA**CTGC**TGAGACCGAATTCTCGCCCT |
| Clone 4 | AGAAGTGAAGCTTGGTCTCA**TCAG**GGGTCATCCAAGAATGTTAT | GCCACCATCTGTTCCTTTAA**CTGC**TGAGACCGAATTCTCGCCCT |
| Clone 5 | AGAAGTGAAGCTTGGTCTCA**TCAG**GGGTCATCCAAGAATGTTAT | GCCACCATCTGTTCCTTTAA**CTGC**TGAGACCGAATTCTCGCCCT |
| **8) MoClo Plant Part A10 (CDS1 module) to GreenGate pGGD000 (Antisense/Sense)**<br>**Colonies:** 142 / **Efficiency**: 100% | | |
| **Vector** | **Position 1** | **Position 2** |
| Original | TTAATCACTCTGTGGTCTCA**AATG**GGGTCATCCAAGAATGTTAT | GCCACCATCTGTTCCTTTAA**GCTT**TGAGACCACGAAGTGGCTCT |
| Desired | AGAAGTGAAGCTTGGTCTCA**TCAG**GGGTCATCCAAGAATGTTAT | GCCACCATCTGTTCCTTTAA**CTGC**TGAGACCGAATTCTCGCCCT |
| Clone 1 | AGAAGTGAAGCTTGGTCTCA**TCAG**GGGTCATCCAAGAATGTTAT | GCCACCATCTGTTCCTTTAA**CTGC**TGAGACCGAATTCTCGCCCT |
| Clone 2 | AGAAGTGAAGCTTGGTCTCA**TCAG**GGGTCATCCAAGAATGTTAT | GCCACCATCTGTTCCTTTAA**CTGC**TGAGACCGAATTCTCGCCCT |
| Clone 3 | AGAAGTGAAGCTTGGTCTCA**TCAG**GGGTCATCCAAGAATGTTAT | GCCACCATCTGTTCCTTTAA**CTGC**TGAGACCGAATTCTCGCCCT |
| Clone 4 | AGAAGTGAAGCTTGGTCTCA**TCAG**GGGTCATCCAAGAATGTTAT | GCCACCATCTGTTCCTTTAA**CTGC**TGAGACCGAATTCTCGCCCT |
| Clone 5 | AGAAGTGAAGCTTGGTCTCA**TCAG**GGGTCATCCAAGAATGTTAT | GCCACCATCTGTTCCTTTAA**CTGC**TGAGACCGAATTCTCGCCCT |
| **9) MoClo Plant Part A12 (3U + Ter module) to GreenGate pGGE000 (Sense/Sense)**<br>**Colonies:** 368 / **Efficiency**: 100% | | |
| **Vector** | **Position 1** | **Position 2** |
| Original | TTAATCACTCTGTGGTCTCA**GCTT**GCTCTCAAGATCAAAGGCTT | ATTTTAAGATCGCACCATTT**CGCT**TGAGACCACGAAGTGGCTCT |
| Desired | AGAAGTGAAGCTTGGTCTCA**CTGC**GCTCTCAAGATCAAAGGCTT | ATTTTAAGATCGCACCATTT**ACTA**TGAGACCGAATTCTCGCCCT |
| Clone 1 | AGAAGTGAAGCTTGGTCTCA**CTGC**GCTCTCAAGATCAAAGGCTT | ATTTTAAGATCGCACCATTT**ACTA**TGAGACCGAATTCTCGCCCT |
| Clone 2 | AGAAGTGAAGCTTGGTCTCA**CTGC**GCTCTCAAGATCAAAGGCTT | ATTTTAAGATCGCACCATTT**ACTA**TGAGACCGAATTCTCGCCCT |
| Clone 3 | AGAAGTGAAGCTTGGTCTCA**CTGC**GCTCTCAAGATCAAAGGCTT | ATTTTAAGATCGCACCATTT**ACTA**TGAGACCGAATTCTCGCCCT |
| Clone 4 | AGAAGTGAAGCTTGGTCTCA**CTGC**GCTCTCAAGATCAAAGGCTT | ATTTTAAGATCGCACCATTT**ACTA**TGAGACCGAATTCTCGCCCT |
| Clone 5 | AGAAGTGAAGCTTGGTCTCA**CTGC**GCTCTCAAGATCAAAGGCTT | ATTTTAAGATCGCACCATTT**ACTA**TGAGACCGAATTCTCGCCCT |
| **10) MoClo Plant Part A12 (3U + Ter module) to GreenGate pGGE000 (Antisense/Antisense)**<br>**Colonies:** 492 / **Efficiency**: 100% | | |
| **Vector** | **Position 1** | **Position 2** |
| Original | TTAATCACTCTGTGGTCTCA**GCTT**GCTCTCAAGATCAAAGGCTT | ATTTTAAGATCGCACCATTT**CGCT**TGAGACCACGAAGTGGCTCT |
| Desired | AGAAGTGAAGCTTGGTCTCA**CTGC**GCTCTCAAGATCAAAGGCTT | ATTTTAAGATCGCACCATTT**ACTA**TGAGACCGAATTCTCGCCCT |
| Clone 1 | AGAAGTGAAGCTTGGTCTCA**CTGC**GCTCTCAAGATCAAAGGCTT | ATTTTAAGATCGCACCATTT**ACTA**TGAGACCGAATTCTCGCCCT |
| Clone 2 | AGAAGTGAAGCTTGGTCTCA**CTGC**GCTCTCAAGATCAAAGGCTT | ATTTTAAGATCGCACCATTT**ACTA**TGAGACCGAATTCTCGCCCT |
| Clone 3 | AGAAGTGAAGCTTGGTCTCA**CTGC**GCTCTCAAGATCAAAGGCTT | ATTTTAAGATCGCACCATTT**ACTA**TGAGACCGAATTCTCGCCCT |
| Clone 4 | AGAAGTGAAGCTTGGTCTCA**CTGC**GCTCTCAAGATCAAAGGCTT | ATTTTAAGATCGCACCATTT**ACTA**TGAGACCGAATTCTCGCCCT |
| Clone 5 | AGAAGTGAAGCTTGGTCTCA**CTGC**GCTCTCAAGATCAAAGGCTT | ATTTTAAGATCGCACCATTT**ACTA**TGAGACCGAATTCTCGCCCT |
| **11) MoClo Plant Part A12 (3U + Ter module) to GreenGate pGGE000 (Sense/Antisense)**<br>**Colonies:** 208 / **Efficiency**: 100% | | |
| **Vector** | **Position 1** | **Position 2** |
| Original | TTAATCACTCTGTGGTCTCA**GCTT**GCTCTCAAGATCAAAGGCTT | ATTTTAAGATCGCACCATTT**CGCT**TGAGACCACGAAGTGGCTCT |

**SUPPLEMENTARY MATERIALS**

| | | |
|---|---|---|
| Desired | AGAAGTGAAGCTTGGTCTCA**CTGC**GCTCTCAAGATCAAAGGCTT | ATTTTAAGATCGCACCATTT**ACTA**TGAGACCGAATTCTCGCCCT |
| Clone 1 | AGAAGTGAAGCTTGGTCTCA**CTGC**GCTCTCAAGATCAAAGGCTT | ATTTTAAGATCGCACCATTT**ACTA**TGAGACCGAATTCTCGCCCT |
| Clone 2 | AGAAGTGAAGCTTGGTCTCA**CTGC**GCTCTCAAGATCAAAGGCTT | ATTTTAAGATCGCACCATTT**ACTA**TGAGACCGAATTCTCGCCCT |
| Clone 3 | AGAAGTGAAGCTTGGTCTCA**CTGC**GCTCTCAAGATCAAAGGCTT | ATTTTAAGATCGCACCATTT**ACTA**TGAGACCGAATTCTCGCCCT |
| Clone 4 | AGAAGTGAAGCTTGGTCTCA**CTGC**GCTCTCAAGATCAAAGGCTT | ATTTTAAGATCGCACCATTT**ACTA**TGAGACCGAATTCTCGCCCT |
| Clone 5 | AGAAGTGAAGCTTGGTCTCA**CTGC**GCTCTCAAGATCAAAGGCTT | ATTTTAAGATCGCACCATTT**ACTA**TGAGACCGAATTCTCGCCCT |
| **12) MoClo Plant Part A12 (3U + Ter module) to GreenGate pGGE000 (Antisense/Sense)**<br>**Colonies:** 471 / **Efficiency**: 100% | | |
| **Vector** | **Position 1** | **Position 2** |
| Original | TTAATCACTCTGTGGTCTCA**GCTT**GCTCTCAAGATCAAAGGCTT | ATTTTAAGATCGCACCATTT**CGCT**TGAGACCACGAAGTGGCTCT |
| Desired | AGAAGTGAAGCTTGGTCTCA**CTGC**GCTCTCAAGATCAAAGGCTT | ATTTTAAGATCGCACCATTT**ACTA**TGAGACCGAATTCTCGCCCT |
| Clone 1 | AGAAGTGAAGCTTGGTCTCA**CTGC**GCTCTCAAGATCAAAGGCTT | ATTTTAAGATCGCACCATTT**ACTA**TGAGACCGAATTCTCGCCCT |
| Clone 2 | AGAAGTGAAGCTTGGTCTCA**CTGC**GCTCTCAAGATCAAAGGCTT | ATTTTAAGATCGCACCATTT**ACTA**TGAGACCGAATTCTCGCCCT |
| Clone 3 | AGAAGTGAAGCTTGGTCTCA**CTGC**GCTCTCAAGATCAAAGGCTT | ATTTTAAGATCGCACCATTT**ACTA**TGAGACCGAATTCTCGCCCT |
| Clone 4 | AGAAGTGAAGCTTGGTCTCA**CTGC**GCTCTCAAGATCAAAGGCTT | ATTTTAAGATCGCACCATTT**ACTA**TGAGACCGAATTCTCGCCCT |
| Clone 5 | AGAAGTGAAGCTTGGTCTCA**CTGC**GCTCTCAAGATCAAAGGCTT | ATTTTAAGATCGCACCATTT**ACTA**TGAGACCGAATTCTCGCCCT |

**SUPPLEMENTARY MATERIALS**

Supplementary Table 2: Clone conversion within standard

Partial sequences of the original sequence, the desired sequence and five sequenced clones are shown in each table. The BsaI recognition motif is underlined. The 4-bp overhangs that are generated by BsaI digestion are shown in green, flanked by 20 bp of sequence at both sides. The sequence highlighted in gray is the backbone sequence. Mutations that deviate from the desired sequence are highlighted in red. The number of colonies and the cloning efficiency is also indicated. Experiment number is indicated in top left corner.

**13) GreenGate pGG-D-GFP*-E to GreenGate pGG-B-GFP-C**
**Colonies**: 88 / **Efficiency**: 80%

| Vector | Position 1 | Position 2 |
|---|---|---|
| Original | AGAAGTGAAGCTTGGTCTCA**TCAG**CTATGGTGAGCAAGGGCGAG | TGGACGAGCTGTACAAGTAA**CTGC**TGAGACCGAATTCTCGCCCT |
| Desired | AGAAGTGAAGCTTGGTCTCA**AACA**CTATGGTGAGCAAGGGCGAG | GCATGGACGAGCTGTACAAG**GGCT**TGAGACCGAATTCTCGCCCT |
| Clone 1 | AGAAGTGAAGCTTGGTCTCA**AACA**CTATGGTGAGCAAGGGCGAG | GCATGGACGAGCTGTACAAG**GGCT**TGAGACCGAATTCTCGCCCT |
| Clone 2 | AGAAGTGAAGCTTGGTCTCA**AACA**CTATGGTGAGCAAGGGCGAG | GCATGGACGAGCTGTACAAG**GGCT**TGAGACCGAATTCTCGCCCT |
| Clone 3 | AGAAGTGAAGCTTGGTCTCA**AACA**CTATGGTGAGCAAGGGCGAG | GCATGGACGAGCTGTACAAG**GGCT**TGAGACCGAATTCTCGCCCT |
| Clone 4 | AGAAGTGAAGCTTGGTCTCA**AACA**CTATGGTGAGCAAGGGCGAG | GCATGGACGAGCTGTACAAG**GGCT**TGAGACCGAATTCTCGCCCT |
| Clone 5 | AGAAGTGAAGCTTGGTCTCA**AACA**CTATGGTGAGCAAGGGCGAG | TGGACGAGCTGTACAAGTAA**CTGC**TGAGACCGAATTCTCGCCCT |

**14) GreenGate pGG-E-MBP*-F to GreenGate pGG-B-MBP-C**
**Colonies**: 327 / **Efficiency**: 80%

| Vector | Position 1 | Position 2 |
|---|---|---|
| Original | AGAAGTGAAGCTTGGTCTCA**CTGC**AAAATCGAAGAAGGTAAACTG | TGAAAGACGCGCAGACTTAG**ACTA**TGAGACCGAATTCTCGCCCT |
| Desired | AGAAGTGAAGCTTGGTCTCA**AACA**GCATGAAAATCGAAGAAGGTAAACTG | CCCTGAAAGACGCGCAGACT**GGCT**TGAGACCGAATTCTCGCCCT |
| Clone 1 | AGAAGTGAAGCTTGGTCTCA**AACA**GCATGAAAATCGAAGAAGGTAAACTG | CCCTGAAAGACGCGCAGACT**GGCT**TGAGACCGAATTCTCGCCCT |
| Clone 2 | AGAAGTGAAGCTTGGTCTCA**AACA**GCATGAAAATCGAAGAAGGTAAACTG | CCCTGAAAGACGCGCAGACT**GGCT**TGAGACCGAATTCTCGCCCT |
| Clone 3 | AGAAGTGAAGCTTGGTCTCA**AACA**GCATGAAAATCGAAGAAGGTAAACTG | CCCTGAAAGACGCGCAGACT**GGCT**TGAGACCGAATTCTCGCCCT |
| Clone 4 | AGAAGTGAAGCTTGGTCTCA**AACA**GCATGAAAATCGAAGAAGGTAAACTG | CCCTGAAAGACGCGCAGACT**GGCT**TGAGACCGAATTCTCGCCCT |
| Clone 5 | AGAAGTGAAGCTTGGTCTCA**AACA**GCATGAAAATCGAAGAAGGTAAACTG | CCCTGAAAGACGCGCAGACT**GGCT**TTAGACCGAATTCTCGCCCT |

**15) GreenGate pGG-B-GST-C to GreenGate pGG-A-GST-B (Gel extraction + pGGC000 backbone)**
**Colonies**: 29 / **Efficiency**: 80%

| Vector | Position 1 | Position 2 |
|---|---|---|
| Original | AGAAGTGAAGCTTGGTCTCA**AACA**CCATGTCCCCTATACTAGGTTA | AATCGGATCTAGTTCCGCGT**GGCT**TGAGACCGAATTCTCGCCCT |
| Desired | AGAAGTGAAGCTTGGTCTCA**ACCT**ATGTCCCCTATACTAGGTTA | AATCGGATCTAGTTCCGCGT**TCAA**CATGAGACCGAATTCTCGCCCT |
| Clone 1 | AGAAGTGAAGCTTGGTCTCA**ACCT**ATGTCCCCTATACTAGGTTA | AATCGGATCTAGTTCCGCGT**TCAA**CATGAGACCGAATTCTCGCCCT |
| Clone 2 | AGAAGTGAAGCTTGGTCTCA**ACCT**ATGTCCCCTATACTAGGTTA | AATCGGATCTAGTTCCGCGT**TCAA**CATGAGACCGAATTCTCGCCCT |
| Clone 3 | AGAAGTGAAGCTTGGTCTCA**ACCT**ATGTCCCCTATACTAGGTTA | AATCGGATCTAGTTCCACGT**TCAA**CATGAGACCGAATTCTCGCCCT |
| Clone 4 | AGAAGTGAAGCTTGGTCTCA**ACCT**ATGTCCCCTATACTAGGTTA | AATCGGATCTAGTTCCGCGT**TCAA**CATGAGACCGAATTCTCGCCCT |
| Clone 5 | AGAAGTGAAGCTTGGTCTCA**ACCT**ATGTCCCCTATACTAGGTTA | AATCGGATCTAGTTCCGCGT**TCAA**CATGAGACCGAATTCTCGCCCT |

SUPPLEMENTARY MATERIALS

#### 16) GreenGate pGG-A-MtU6-B to GreenGate pGG-B-MtU6-C

**Colonies**: 27 / **Efficiency**: 20%

| Vector | Position 1 | Position 2 |
|---|---|---|
| Original | AGAAGTGAAGCTTGGTCTCA**ACCT**ATGCCTATCTTATATGATCA | CTTGTACAAAGTTGGCATTA**AACA**TGAGACCGAATTCTCGCCCT |
| Desired | AGAAGTGAAGCTTGGTCTCA**AACA**ATGCCTATCTTATATGATCA | CTTGTACAAAGTTGGCATTA**GGCT**TGAGACCGAATTCTCGCCCT |
| Clone 1 | AGAAGTGAAGCTTGGTCTCA**ACCT**ATGCCTATCTTATATGATCA | CTTGTACAAAGTTGGCATTA**AACA**TGAGACCGAATTCTCGCCCT |
| Clone 2 | AGAAGTGAAGCTTGGTCTCA**AACA**ATGCCTATCTTATATGATCA | CTTGTACAAAGTTGGCATTA**GGCT**TGAGACCGAATTCTCGCCCT |
| Clone 3 | Mix of AACA and ACCT | Mix of AACA and ACCT |
| Clone 4 | AGAAGTGAAGCTTGGTCTCA**ACCT**ATGCCTATCTTATATGATCA | CTTGTACAAAGTTGGCATTA**GGCT**TGAGACCGAATTCTCGCCCT |
| Clone 5 | AGAAGTGAAGCTTGGTCTCA**ACCT**ATGCCTATCTTATATGATCA | CTTGTACAAAGTTGGCATTA**AACA**TGAGACCGAATTCTCGCCCT |

#### 17) GreenGate pGG-A-MtU6-B to GreenGate pGG-B-MtU6-C (pGGC000 GmR)

**Colonies**: 39 / **Efficiency**: 40%

| Vector | Position 1 | Position 2 |
|---|---|---|
| Original | AGAAGTGAAGCTTGGTCTCA**ACCT**ATGCCTATCTTATATGATCA | CTTGTACAAAGTTGGCATTA**AACA**TGAGACCGAATTCTCGCCCT |
| Desired | AGAAGTGAAGCTTGGTCTCA**AACA**ATGCCTATCTTATATGATCA | CTTGTACAAAGTTGGCATTA**GGCT**TGAGACCGAATTCTCGCCCT |
| Clone 1 | AGAAGTGAAGCTTGGTCTCA**AACA**ATGCCTATCTTATATGATCA | CTTGTACAAAGTTGGCATTA**GGCT**TGAGACCGAATTCTCGCCCT |
| Clone 2 | AGAAGTGAAGCTTGGTCT-A**AACA**ATGCCTATCTTATATGATCA | CTTGTACAAAGTTGGCATTA**GGCT**TGAGACCGAATTCTCGCCCT |
| Clone 3 | AGAAGTGAAGCTTGGTCTCA**AACA**ATGCCTATCTTATATGATCA | CTTGTACAAAGTTGGCATTA**GGCT**TGAGACCGAATTCTCGCCCT |
| Clone 4 | AGAAGTGAAGCTTGGTCTCA**ACCT**ATGCCTATCTTATATGATCA | CTTGTACAAAGTTGGCATTA**AACA**TGAGACCGAATTCTCGCCCT |
| Clone 5 | AGAAGTGAAGCTTGGTCT-A**AACA**ATGCCTATCTTATATGATCA | CTTGTACAAAGTTGGCATTA**GGCT**TGAGACCGAATTCTCGCCCT |

#### 18) GreenGate pGG-A-MtU6-B to GreenGate pGG-C-MtU6-D

**Colonies**: 28 / **Efficiency**: 40%

| Vector | Position 1 | Position 2 |
|---|---|---|
| Original | AGAAGTGAAGCTTGGTCTCA**ACCT**ATGCCTATCTTATATGATCA | CTTGTACAAAGTTGGCATTA**AACA**TGAGACCGAATTCTCGCCCT |
| Desired | AGAAGTGAAGCTTGGTCTCA**GGCT**ATGCCTATCTTATATGATCA | CTTGTACAAAGTTGGCATTA**TCAG**TGAGACCGAATTCTCGCCCT |
| Clone 1 | AGAAGTGAAGCTTGGTCTCA**GGCT**ATGCCTATCTTATATGATCA | CTTGTACAAAGTTGGCATTA**TCAG**TGAGACCGAATTCTCGCCCT |
| Clone 2 | AGAAGTGAAGCTTGGTCTCA**ACCT**ATGCCTATCTTATATGATCA | CTTGTACAAAGTTGGCATTA**AACA**TGAGACCGAATTCTCGCCCT |
| Clone 3 | AGAAGTGAAGCTTGGTCTCA**ACCT**ATGCCTATCTTATATGATCA | CTTGTACAAAGTTGGCATTA**AACA**TGAGACCGAATTCTCGCCCT |
| Clone 4 | AGAAGTGAAGCTTGGTCTCA**GGCT**ATGCCTATCTTATATGATCA | CTTGTACAAAGTTGGCATTA**TCAG**TGAGACCGAATTCTCGCCCT |
| Clone 5 | AGAAGTGAAGCTTGGTCTCA**ACCT**ATGCCTATCTTATATGATCA | CTTGTACAAAGTTGGCATTA**AACA**TGAGACCGAATTCTCGCCCT |

#### 19) GreenGate pGG-A-MtU6-B to GreenGate pGG-C-MtU6-D (pGGC000 GmR)

**Colonies**: 72 / **Efficiency**: 40%

| Vector | Position 1 | Position 2 |
|---|---|---|
| Original | AGAAGTGAAGCTTGGTCTCA**ACCT**ATGCCTATCTTATATGATCA | CTTGTACAAAGTTGGCATTA**AACA**TGAGACCGAATTCTCGCCCT |
| Desired | AGAAGTGAAGCTTGGTCTCA**GGCT**ATGCCTATCTTATATGATCA | CTTGTACAAAGTTGGCATTA**TCAG**TGAGACCGAATTCTCGCCCT |
| Clone 1 | AGAAGTGAAGCTTGGTCTCA**GGCT**ATGCCTATCTTATATGATCA | CTTGTACAAAGTTGGCATTA**TCAG**TG-GACCGAATTCTCGCCCT |
| Clone 2 | AGAAGTGAAGCTTGGTCTCA**GGCT**ATGCCTATCTTATATGATCA | CTTGTACAAAGTTGGCATTA**TCAG**TGAGACCGAATTCTCGCCCT |
| Clone 3 | AGAAGTGAAGCTTGGTCTCA**GGCT**ATGCCTATCTTATATGATCA | CTTGTACAAAGTTGGCATTA**TCAG**TGAGACCGAATTCTCGCCCT |

### SUPPLEMENTARY MATERIALS

| | | |
|---|---|---|
| Clone 4 | AGAAGTGAAGCTTGTTCTCA**GGCT**ATGCCTATCTTATATGATCA | CTTGTACAAAGTTGGCATTA**TCAG**TGAGACCGAATTCTCGCCCT |
| Clone 5 | AGAAGTGAAGCTTGGTCTCA**ACCT**ATGCCTATCTTATATGATCA | CTTGTACAAAGTTGGCATTA**AACA**TGAGACCGAATTCTCGCCCT |

**SUPPLEMENTARY MATERIALS**

Supplementary Table 3: Replacement of BsaI recognition site with other TypeIIS site

Partial sequences of the original sequence, the desired sequence and five sequenced clones are shown in each table. The positions in the table refer to the junctions in final tagged clone, for the original untagged vector the end that is tagged is shown twice in the table. The BsaI/AarI/SapI recognition motifs are underlined. The overhangs that are generated the restriction enzymes are shown in bold. The sequence highlighted in gray is the backbone sequence. Mutations that deviate from the desired sequence are highlighted in red. The number of colonies and the cloning efficiency is also indicated. Experiment number is indicated in top left corner.

**20) MoClo Plant Part A10 (CDS1 module) to GreenGate (AarI overhangs)**
**Colonies:** 750 / **Efficiency**: 100%

| Vector | Position 1 | Position 2 |
|---|---|---|
| Original | TGGCCGATTCATTAATCACTCTGTGGTCTCA**AATG**GGGTCATCCAAGAATGTTAT | GCCACCATCTGTTCCTTTAA**GCTT**TGAGACCACGAAGTGGCTCTTCAGTGGACGA |
| Desired | ACACTATAGAAGTGAAGCTTCACCTGCAATA**TCAG**GGGTCATCCAAGAATGTTAT | GCCACCATCTGTTCCTTTAA**CTGC**TCGTGCAGGTGGAATTCTCGCCCTATAGTGA |
| Clone 1 | ACACTATAGAAGTGAAGCTTCACCTGCAATA**TCAG**GGGTCATCCAAGAATGTTAT | GCCACCATCTGTTCCTTTAA**CTGC**TCGTGCAGGTGGAATTCTCGCCCTATAGTGA |
| Clone 2 | ACACTATAGAAGTGAAGCTTCACCTGCAATA**TCAG**GGGTCATCCAAGAATGTTAT | GCCACCATCTGTTCCTTTAA**CTGC**TCGTGCAGGTGGAATTCTCGCCCTATAGTGA |
| Clone 3 | ACACTATAGAAGTGAAGCTTCACCTGCAATA**TCAG**GGGTCATCCAAGAATGTTAT | GCCACCATCTGTTCCTTTAA**CTGC**TCGTGCAGGTGGAATTCTCGCCCTATAGTGA |
| Clone 4 | ACACTATAGAAGTGAAGCTTCACCTGCAATA**TCAG**GGGTCATCCAAGAATGTTAT | GCCACCATCTGTTCCTTTAA**CTGC**TCGTGCAGGTGGAATTCTCGCCCTATAGTGA |
| Clone 5 | ACACTATAGAAGTGAAGCTTCACCTGCAATA**TCAG**GGGTCATCCAAGAATGTTAT | GCCACCATCTGTTCCTTTAA**CTGC**TCGTGCAGGTGGAATTCTCGCCCTATAGTGA |

**21) MoClo Plant Part A12 (3U + Ter module) to GreenGate (SapI overhangs)**
**Colonies:** 1520 / **Efficiency**: 80%

| Vector | Position 1 | Position 2 |
|---|---|---|
| Original | CGATTCATTAATCACTCTGTGGTCTCA**GCTT**GCTCTCAAGATCAAAGGCTT | ATTTTAAGATCGCACCATTT**CGCT**TGAGACCACGAAGTGGCTCTTCAGTGG |
| Desired | ACACTATAGAAGTGAAGCTTGCTCTTCA**TGA**GCTCTCAAGATCAAAGGCTT | ATTTTAAGATCGCACCATTT**GAT**TGAAGAGCGAATTCTCGCCCTATAGTGA |
| Clone 1 | ACACTATAGAAGTGAAGCTTGGTCTTCA**TGA**GCTCTCAAGATCAAAGGCTT | ATTTTAAGATCGCACCATTT**GAT**TGAAGAGCGAATTCTCGCCCTATAGTGA |
| Clone 2 | ACACTATAGAAGTGAAGCTTGCTCTTCA**TGA**GCTCTCAAGATCAAAGGCTT | ATTTTAAGATCGCACCATTT**GAT**TGAAGAGCGAATTCTCGCCCTATAGTGA |
| Clone 3 | ACACTATAGAAGTGAAGCTTGCTCTTCA**TGA**GCTCTCAAGATCAAAGGCTT | ATTTTAAGATCGCACCATTT**GAT**TGAAGAGCGAATTCTCGCCCTATAGTGA |
| Clone 4 | ACACTATAGAAGTGAAGCTTGCTCTTCA**TGA**GCTCTCAAGATCAAAGGCTT | ATTTTAAGATCGCACCATTT**GAT**TGAAGAGCGAATTCTCGCCCTATAGTGA |
| Clone 5 | ACACTATAGAAGTGAAGCTTGCTCTTCA**TGA**GCTCTCAAGATCAAAGGCTT | ATTTTAAGATCGCACCATTT**GAT**TGAAGAGCGAATTCTCGCCCTATAGTGA |

**SUPPLEMENTARY MATERIALS**

Supplementary Table 4: Clone tagging with PCR products or donor plasmids

Partial sequences of the original sequence, the desired sequence and five sequenced clones are shown in each table. The positions in the table refer to the junctions in final tagged clone, for the original untagged vector the end that is tagged is shown twice in the table. The BsaI recognition motif is underlined. The 4-bp overhangs that are generated by BsaI digestion are shown in bold. The sequence that is tagged is highlighted in blue, the tag in green. The sequence highlighted in gray is the backbone sequence. The sequence highlighted in yellow is a linker that was included in the oligo. Mutations that deviate from the desired sequence are highlighted in red. The number of colonies and the cloning efficiency is also indicated. Experiment number is indicated in top left corner.

**24) GUS C-terminal GFP-fusion (PCR product) to pGGC0000 (CbR)**
**Colonies:** 184 / **Efficiency**: 80%

| Vector | Position 1 | Position 2 | Position 3 |
|---|---|---|---|
| Original | AGAAGTGAAGCTTGGTCTCA**GGCT**CAACAATGGTCCGTCCTGTA | CGCAGCAGGGAGGCAAACAA**TCAG**TGAGACCGAATTCTCGCCCT | CGCAGCAGGGAGGCAAACAA**TCAG**TGAGACCGAATTCTCGCCCT |
| Desired | AGAAGTGAAGCTTGGTCTCA**GGCT**CAACAATGGTCCGTCCTGTA | GGGAGGCAAACAA**GGTGGATCAGGCGGAAGT**ATGGTGAGCAAGG | GCATGGACGAGCTGTACAAG**TCAG**TGAGACCGAATTCTCGCCCT |
| Clone 1 | AGAAGTGAAGCTTGGTCTCA**GGCT**CAACAATGGTCCGTCCTGTA | GGGAGGCAAACAA**GGTGGATCAGGCGGAAGT**ATGGTGAGCAAGG | GCATGGACGAGCTGTACAAG**TCAG**TGAGACCGAATTCTCGCCCT |
| Clone 2 | GAAGTGAAGCTTGGGTCTCA**GGCT**CAACAATGGTCCGTCCTGTA | GGGAGGCAAACAA**GGTGGATCAGGCGGAAGT**ATGGTGAGCAAGG | GCATGGACGAGCTGTACAAG**TCAG**TGAGACCGAATTCTCGCCCT |
| Clone 3 | AGAAGTGAAGCTTGGTCTCA**GGCT**CAACAATGGTCCGTCCTGTA | GGGAGGCAAACAA**GGTGGATCAGGCGGAAGT**ATGGTGAGCAAGG | GCATGGACGAGCTGTACAAG**TCAG**TGAGACCGAATTCTCGCCCT |
| Clone 4 | AGAAGTGAAGCTTGGTCTCA**GGCT**CAACAATGGTCCGTCCTGTA | GGGAGGCAAACAA**GGTGGATCAGGCGGAAGT**ATGGTGAGCAAGG | GCATGGACGAGCTGTACAAG**TCAG**TGAGACCGAATTCTCGCCCT |
| Clone 5 | AGAAGTGAAGCTTGGTCTCA**GGCT**CAACAATGGTCCGTCCTGTA | GGGAGGCAAACAA**GGTGGATCAGGCGGAAGT**ATGGTGAGCAAGG | GCATGGACGAGCTGTACAAG**TCAG**TGAGACCGAATTCTCGCCCT |

**25) Integrase 4 C-terminal V5-tag-fusion (donor vector) to pGGC0000 (CbR)**
**Colonies:** 94 / **Efficiency**: 60%

| Vector | Position 1 | Position 2 | Position 3 |
|---|---|---|---|
| Original | AGAAGTGAAGCTTGGTCTCA**GGCT**CCATGATTACGACCAGAAAG | TAGATATACATTGGACCTTT**TCAG**TGAGACCGAATTCTCGCCCT | TAGATATACATTGGACCTTT**TCAG**TGAGACCGAATTCTCGCCCT |
| Desired | AGAAGTGAAGCTTGGTCTCA**GGCT**CCATGATTACGACCAGAAAG | TAGATATACATTGGACCTTTGGTAAGCCAATCCCTAATCCTCTC | TCGGACTCGACTCAACCTAA**TCAG**TGAGACCGAATTCTCGCCCT |
| Clone 1 | AGAAGTGAAGCTTGGTCTCA**GGCT**CCATGATTACGACCAGAAAG | TAGATATACATTGGACCTTT**TCAG**TGAGACCGAATTCTCGCCCT | TAGATATACATTGGACCTTT**TCAG**TGAGACCGAATTCTCGCCCT |
| Clone 2 | AGAAGTGAAGCTTGGTCTCA**GGCT**CCATGATTACGACCAGAAAG | TAGATATACATTGGACCTTTGGTAAGCCAATCCCTAATCCTCTC | TCGGACTCGACTCAACCTAA**TCAG**TGAGACCGAATTCTCGCCCT |
| Clone 3 | AGAAGTGAAGCTTGGTCTCA**GGCT**CCATGATTACGACCAGAAAG | TAGATATACATTGGACCTTTGGTAAGCCAATCCCTAATCCTCTC | TCGGACTCGACTCAACCTAA**TCAG**TGAGACCGAATTCTCGCCCT |
| Clone 4 | AGAAGTGAAGCTTGGTCTCA**GGCT**CCATGATTACGACCAGAAAG | TAGATATACATTGGACCTTTGGTAAGCCAATCCCTAATCCTCTC | TCGGACTCGACTCAACCTAA**TCAG**TGAGACCGAATTCTCGCCCT |
| Clone 5 | AGAAGTGAAGCTTGGTCTCA**GGCT**CCATGATTACGACCAGAAAG | TAGATATACATTGGACCTTT**TCAG**TGAGACCGAATTCTCGCCCT | TAGATATACATTGGACCTTT**TCAG**TGAGACCGAATTCTCGCCCT |

**26) Csy4 N-terminal mCherry-fusion (donor vector) to pGGC0000 (CbR)**
**Colonies:** 29 / **Efficiency**: 0%

| Vector | Position 1 | Position 2 | Position 3 |
|---|---|---|---|
| Original | AGAAGTGAAGCTTGGTCTCA**AACA**ATGGATCATTATCTTGATAT | AGAAGTGAAGCTTGGTCTCA**AACA**ATGGATCATTATCTTGATAT | TTGAAGAAAATCCTGGACCC**GGCT**TGAGACCGAATTCTCGCCCT |
| Desired | AGAAGTGAAGCTTGGTCTCA**AACA**ATGGTGAGCAAGGGCGAGGA | GACGAGCTGTACAAG**GGAGGTTCA**ATGGATCATTATCTTGATAT | TTGAAGAAAATCCTGGACCC**GGCT**TGAGACCGAATTCTCGCCCT |
| Clone 1 | AAGTGAAGCTTGGTCTCA**GGCTCC**ATGGTGAGCAAGGGCGAGGA | AAGTGAAGCTTGGTCTCA**GGCTCC**ATGGTGAGCAAGGGCGAGGA | GCATGGACGAGCTGTACAAG**TCAG**TGAGACCGAATTCTCGCCCT |
| Clone 2 | AAGTGAAGCTTGGTCTCA**GGCTCC**ATGGTGAGCAAGGGCGAGGA | AAGTGAAGCTTGGTCTCA**GGCTCC**ATGGTGAGCAAGGGCGAGGA | GCATGGACGAGCTGTACAAG**TCAG**TGAGACCGAATTCTCGCCCT |
| Clone 3 | AAGTGAAGCTTGGTCTCA**GGCTCC**ATGGTGAGCAAGGGCGAGGA | AAGTGAAGCTTGGTCTCA**GGCTCC**ATGGTGAGCAAGGGCGAGGA | GCATGGACGAGCTGTACAAG**TCAG**TGAGACCGAATTCTCGCCCT |

### SUPPLEMENTARY MATERIALS

| | | | |
|---|---|---|---|
| Clone 4 | AAGTGAAGCTTGGTCTCAGGCTCCATGGTGAGCAAGGGCGAGGA | AAGTGAAGCTTGGTCTCAGGCTCCATGGTGAGCAAGGGCGAGGA | GCATGGACGAGCTGTACAAGTCAGTGAGACCGAATTCTCGCCCT |
| Clone 5 | AAGTGAAGCTTGGTCTCAGGCTCCATGGTGAGCAAGGGCGAGGA | AAGTGAAGCTTGGTCTCAGGCTCCATGGTGAGCAAGGGCGAGGA | GCATGGACGAGCTGTACAAGTCAGTGAGACCGAATTCTCGCCCT |

**27) GUS C-terminal GFP-fusion (PCR product) to pGGC0000 (GmR)**

**Colonies:** 112/ **Efficiency**: 80%

| Vector | Position 1 | Position 2 | Position 3 |
|---|---|---|---|
| Original | AGAAGTGAAGCTTGGTCTCAGGCTCAACAATGGTCCGTCCTGTA | CGCAGCAGGGAGGCAAACAATCAGTGAGACCGAATTCTCGCCCT | CGCAGCAGGGAGGCAAACAATCAGTGAGACCGAATTCTCGCCCT |
| Desired | AGAAGTGAAGCTTGGTCTCAGGCTCAACAATGGTCCGTCCTGTA | GGGAGGCAAACAAGGTGGATCAGGCGGAAGTATGGTGAGCAAGG | GCATGGACGAGCTGTACAAGTCAGTGAGACCGAATTCTCGCCCT |
| Clone 1 | AGAAGTGAAGCTTGGTCTCAGGCTCAACAATGGTCCGTCCTGTA | GGGAGGCAAACAAGGTGGATCAGGCGGAAGTATGGTGAGCAAGG | GCATGGACGAGCTGTACAAGTCAGTGAGACCGAATTCTCGCCCT |
| Clone 2 | AGAAGTGAAGCTTGGTCTCAGGCTCAACAATGGTCCGTCCTGTA | GGGAGGCAAACAAGGTGGATCAGGCGGAAGTATGGTGAGCAAGG | GCATGGACGAGCTGTACAAGT-AGTGAGACCGAATTCTCGCCCT |
| Clone 3 | AGAAGTGAAGCTTGGTCTCAGGCTCAACAATGGTCCGTCCTGTA | GGGAGGCAAACAAGGTGGATCAGGCGGAAGTATGGTGAGCAAGG | GCATGGACGAGCTGTACAAGTCAGTGAGACCGAATTCTCGCCCT |
| Clone 4 | AGAAGTGAAGCTTGGTCTCAGGCTCAACAATGGTCCGTCCTGTA | GGGAGGCAAACAAGGTGGATCAGGCGGAAGTATGGTGAGCAAGG | GCATGGACGAGCTGTACAAGTCAGTGAGACCGAATTCTCGCCCT |
| Clone 5 | AGAAGTGAAGCTTGGTCTCAGGCTCAACAATGGTCCGTCCTGTA | GGGAGGCAAACAAGGTGGATCAGGCGGAAGTATGGTGAGCAAGG | GCATGGACGAGCTGTACAAGTCAGTGAGACCGAATTCTCGCCCT |

**28) Integrase 4 C-terminal V5-tag-fusion (vector) to pGGC0000 (GmR)**

**Colonies:** 12 / **Efficiency**: 100%

| Vector | Position 1 | Position 2 | Position 3 |
|---|---|---|---|
| Original | AGAAGTGAAGCTTGGTCTCAGGCTCCATGATTACGACCAGAAAG | TAGATATACATTGGACCTTTTCAGTGAGACCGAATTCTCGCCCT | TAGATATACATTGGACCTTTTCAGTGAGACCGAATTCTCGCCCT |
| Desired | AGAAGTGAAGCTTGGTCTCAGGCTCCATGATTACGACCAGAAAG | TAGATATACATTGGACCTTTGGTAAGCCAATCCCTAATCCTCTC | TCGGACTCGACTCAACCTAATCAGTGAGACCGAATTCTCGCCCT |
| Clone 1 | AGAAGTGAAGCTTGGTCTCAGGCTCCATGATTACGACCAGAAAG | TAGATATACATTGGACCTTTGGTAAGCCAATCCCTAATCCTCTC | TCGGACTCGACTCAACCTAATCAGTGAGACCGAATTCTCGCCCT |
| Clone 2 | AGAAGTGAAGCTTGGTCTCAGGCTCCATGATTACGACCAGAAAG | TAGATATACATTGGACCTTTGGTAAGCCAATCCCTAATCCTCTC | TCGGACTCGACTCAACCTAATCAGTGAGACCGAATTCTCGCCCT |
| Clone 3 | AGAAGTGAAGCTTGGTCTCAGGCTCCATGATTACGACCAGAAAG | TAGATATACATTGGACCTTTGGTAAGCCAATCCCTAATCCTCTC | TCGGACTCGACTCAACCTAATCAGTGAGACCGAATTCTCGCCCT |
| Clone 4 | AGAAGTGAAGCTTGGTCTCAGGCTCCATGATTACGACCAGAAAG | TAGATATACATTGGACCTTTGGTAAGCCAATCCCTAATCCTCTC | TCGGACTCGACTCAACCTAATCAGTGAGACCGAATTCTCGCCCT |
| Clone 5 | AGAAGTGAAGCTTGGTCTCAGGCTCCATGATTACGACCAGAAAG | TAGATATACATTGGACCTTTGGTAAGCCAATCCCTAATCCTCTC | TCGGACTCGACTCAACCTAATCAGTGAGACCGAATTCTCGCCCT |

**29) Csy4 N-terminal mCherry-fusion (vector) to pGGC0000 (GmR)**

**Colonies:** 29 / **Efficiency**: 100%

| Vector | Position 1 | Position 2 | Position 3 |
|---|---|---|---|
| Original | AGAAGTGAAGCTTGGTCTCAAACAATGGATCATTATCTTGATAT | AGAAGTGAAGCTTGGTCTCAAACAATGGATCATTATCTTGATAT | TTGAAGAAAATCCTGGACCCGGCTTGAGACCGAATTCTCGCCCT |
| Desired | AGAAGTGAAGCTTGGTCTCAAACAATGGTGAGCAAGGGCGAGGA | GACGAGCTGTACAAGGGAGGTTCAATGGATCATTATCTTGATAT | TTGAAGAAAATCCTGGACCCGGCTTGAGACCGAATTCTCGCCCT |
| Clone 1 | AGAAGTGAAGCTTGGTCTCAAACAATGGTGAGCAAGGGCGAGGA | GACGAGCTGTACAAGGGAGGTTCAATGGATCATTATCTTGATAT | TTGAAGAAAATCCTGGACCCGGCTTGAGACCGAATTCTCGCCCT |
| Clone 2 | AGAAGTGAAGCTTGGTCTCAAACAATGGTGAGCAAGGGCGAGGA | GACGAGCTGTACAAGGGAGGTTCAATGGATCATTATCTTGATAT | TTGAAGAAAATCCTGGACCCGGCTTGAGACCGAATTCTCGCCCT |
| Clone 3 | AGAAGTGAAGCTTGGTCTCAAACAATGGTGAGCAAGGGCGAGGA | GACGAGCTGTACAAGGGAGGTTCAATGGATCATTATCTTGATAT | TTGAAGAAAATCCTGGACCCGGCTTGAGACCGAATTCTCGCCCT |
| Clone 4 | AGAAGTGAAGCTTGGTCTCAAACAATGGTGAGCAAGGGCGAGGA | GACGAGCTGTACAAGGGAGGTTCAATGGATCATTATCTTGATAT | TTGAAGAAAATCCTGGACCCGGCTTGAGACCGAATTCTCGCCCT |
| Clone 5 | AGAAGTGAAGCTTGGTCTCAAACAATGGTGAGCAAGGGCGAGGA | GACGAGCTGTACAAGGGAGGTTCAATGGATCATTATCTTGATAT | TTGAAGAAAATCCTGGACCCGGCTTGAGACCGAATTCTCGCCCT |

**SUPPLEMENTARY MATERIALS**

Supplementary Table 5: One PCR product to many entries

Partial sequences of the original sequence, the desired sequence and five sequenced clones are shown in each table. The BsaI recognition motif is underlined. The 4-bp overhangs that are generated by BsaI digestion are shown in green, flanked by 20 bp of sequence at both sides. The sequence highlighted in gray is the backbone sequence. Mutations that deviate from the desired sequence are highlighted in red. The number of colonies and the cloning efficiency is also indicated. Experiment number is indicated in top left corner.

| | | |
|---|---|---|
| **30) pGG-A-GST-B (from pEN-R2-GST-L3)** **Colonies**: 336 / **Efficiency**: 100% | | |
| **Vector** | **Position 1** | **Position 2** |
| Original | TACAAAGTGGGCGGAGGTGGCAGCATGTCCCCTATACTAGGTTA | AATCGGATCTAGTTCCGCGTTAAGCAACTTTATTATACAAAGTTGG |
| Desired | AGAAGTGAAGCTTGGTCTCAACCTATGTCCCCTATACTAGGTTA | AATCGGATCTAGTTCCGCGTTCAACATGAGACCGAATTCTCGCCCT |
| Clone 1 | AGAAGTGAAGCTTGGTCTCAACCTATGTCCCCTATACTAGGTTA | AATCGGATCTAGTTCCGCGTTCAACATGAGACCGAATTCTCGCCCT |
| Clone 2 | AGAAGTGAAGCTTGGTCTCAACCTATGTCCCCTATACTAGGTTA | AATCGGATCTAGTTCCGCGTTCAACATGAGACCGAATTCTCGCCCT |
| Clone 3 | AGAAGTGAAGCTTGGTCTCAACCTATGTCCCCTATACTAGGTTA | AATCGGATCTAGTTCCGCGTTCAACATGAGACCGAATTCTCGCCCT |
| Clone 4 | AGAAGTGAAGCTTGGTCTCAACCTATGTCCCCTATACTAGGTTA | AATCGGATCTAGTTCCGCGTTCAACATGAGACCGAATTCTCGCCCT |
| Clone 5 | AGAAGTGAAGCTTGGTCTCAACCTATGTCCCCTATACTAGGTTA | AATCGGATCTAGTTCCGCGTTCAACATGAGACCGAATTCTCGCCCT |
| **31) pGG-B-GST-C (from pEN-R2-GST-L3)** **Colonies**: 410 / **Efficiency**: 100% | | |
| **Vector** | **Position 1** | **Position 2** |
| Original | TACAAAGTGGGCGGAGGTGGCAGCATGTCCCCTATACTAGGTTA | AATCGGATCTAGTTCCGCGTTAAGCAACTTTATTATACAAAGTT |
| Desired | AGAAGTGAAGCTTGGTCTCAAACAATGTCCCCTATACTAGGTTA | AATCGGATCTAGTTCCGCGTGGCTTGAGACCGAATTCTCGCCCT |
| Clone 1 | AGAAGTGAAGCTTGGTCTCAAACAATGTCCCCTATACTAGGTTA | AATCGGATCTAGTTCCGCGTGGCTTGAGACCGAATTCTCGCCCT |
| Clone 2 | AGAAGTGAAGCTTGGTCTCAAACAATGTCCCCTATACTAGGTTA | AATCGGATCTAGTTCCGCGTGGCTTGAGACCGAATTCTCGCCCT |
| Clone 3 | AGAAGTGAAGCTTGGTCTCAAACAATGTCCCCTATACTAGGTTA | AATCGGATCTAGTTCCGCGTGGCTTGAGACCGAATTCTCGCCCT |
| Clone 4 | AGAAGTGAAGCTTGGTCTCAAACAATGTCCCCTATACTAGGTTA | AATCGGATCTAGTTCCGCGTGGCTTGAGACCGAATTCTCGCCCT |
| Clone 5 | AGAAGTGAAGCTTGGTCTCAAACAATGTCCCCTATACTAGGTTA | AATCGGATCTAGTTCCGCGTGGCTTGAGACCGAATTCTCGCCCT |
| **32) pGG-C-GST-D (from pEN-R2-GST-L3)** **Colonies**: 840 / **Efficiency**: 80% | | |
| **Vector** | **Position 1** | **Position 2** |
| Original | TACAAAGTGGGCGGAGGTGGCAGCATGTCCCCTATACTAGGTTA | AATCGGATCTAGTTCCGCGTTAAGCAACTTTATTATACAAAGTT |
| Desired | AGAAGTGAAGCTTGGTCTCAGGCTCTATGTCCCCTATACTAGGTTA | AATCGGATCTAGTTCCGCGTTCAGTGAGACCGAATTCTCGCCCT |
| Clone 1 | AGAAGTGAAGCTTGGTCTCAGGCTCTATGTCCCCTATACTAGGTTA | AATCGGATCTAGTTCCGCGTTCAGTGAGACCGAATTCTCGCCCT |
| Clone 2 | AGAAGTGAAGCTTGGTCTCAGGCTCTATGTCCCCTATACTAGGTTA | AATCGGATCTAGTTCCGCGTTCAGTGAGACCGAATTCTCGCCCT |
| Clone 3 | AGAAGTGAAGCTTGGTCTCAGGCTCTATGTCCCCTATACTAGGTTA | AATCGGATCTAGTTCCGCGTTCAGTGAGACCGAATTCTCGCCCT |
| Clone 4 | AGAAGTGAAGCTTGGTCTCAGGCTCTATGTCCCCTATACTAGGTTA | AATCGGAT-TAGTTCCGCGTTCAGTGAGACCGAATTCTCGCCCT |
| Clone 5 | AGAAGTGAAGCTTGGTCTCAGGCTCTATGTCCCCTATACTAGGTTA | AATCGGATCTAGTTCCGCGTTCAGTGAGACCGAATTCTCGCCCT |

**SUPPLEMENTARY MATERIALS**

**33) pGG-D-GST-E (from pEN-R2-GST-L3)**
**Colonies**: 676 / **Efficiency**: 80%

| Vector | Position 1 | Position 2 |
|---|---|---|
| Original | TACAAAGTGGGCGGAGGTGGCAGCATGTCCCCTATACTAGGTTA | AATCGGATCTAGTTCCGCGTTAAGCAACTTTATTATACAAAGTT |
| Desired | AGAAGTGAAGCTTGGTCTCATCAGCTATGTCCCCTATACTAGGTTA | AATCGGATCTAGTTCCGCGTCTGCTGAGACCGAATTCTCGCCCT |
| Clone 1 | AGAAGTGAAGCTTGGTCTCATCAGCTATGTCCCCTATACTAGGTTA | AATCGGATCTAGTTCCGCGTCTGCTGAGACCGAATTCTCGCCCT |
| Clone 2 | AGAAGTGAAGCTTGGTCTCATCAGCTATGTCCCCTATACTAGGTTA | AATCGGATCTAGTTCCGCGTCTGCTGAGACCGAATTCTCGCCCT |
| Clone 3 | AGAAGTGAAGCTTGGTCTCATCAGCTATGTCCCCTATACTAGGTTA | AATCGGATCTAGT-CCGC-TCTGCTGAGACCGAATTCTCGCCCT |
| Clone 4 | AGAAGTGAAGCTTGGTCTCATCAGCTATGTCCCCTATACTAGGTTA | AATCGGATCTAGTTCCGCGTCTGCTGAGACCGAATTCTCGCCCT |
| Clone 5 | AGAAGTGAAGCTTGGTCTCATCAGCTATGTCCCCTATACTAGGTTA | AATCGGATCTAGTTCCGCGTCTGCTGAGACCGAATTCTCGCCCT |

**34) pGG-E-GST-F (from pEN-R2-GST-L3)**
**Colonies**: 346 / **Efficiency**: 60%

| Vector | Position 1 | Position 2 |
|---|---|---|
| Original | TACAAAGTGGGCGGAGGTGGCAGCATGTCCCCTATACTAGGTTA | AATCGGATCTAGTTCCGCGTTAAGCAACTTTATTATACAAAGTTGG |
| Desired | AGAAGTGAAGCTTGGTCTCACTGCCTATGTCCCCTATACTAGGTTA | AATCGGATCTAGTTCCGCGTTGAACTATGAGACCGAATTCTCGCCC |
| Clone 1 | AGAAGTGAAGCTTGGTCTCACTGCCTATGTCCCCTATACTAGGTTA | AATCGGATCTAGTTCCGCGTTGAACTATGAGACCGAATTCTCGCCC |
| Clone 2 | AGAAGTGAAGCTTGGTCTCACTGCCTATGTCCCCTATACTAGGTTA | AATCGGATCTAGTTCCGCGTTGAACTATGAGACCGAATTCTCGCCC |
| Clone 3 | AGAAGTGAAGCTTGGTCTCACTGC-TATGTCCCCTATACTAGGTTA | AATCGGATCTAGTTCCGCGTTGAACTATGAGACCGAATTCTCGCCC |
| Clone 4 | AGAAGTGAAGCTTGGTCTCACTGCCTATGTCCCCTATACTAGGTTA | AATCGGATCTAGTTCCGCGTTGAACTATGAGACCGAATTCTCGCCC |
| Clone 5 | AGAAGTGAAGCTTGGTCTCACTGCCTATGTCCCCTATACTAGGTTA | AATCGGATCTAGTTCCGCGTTAAGCAACTTTATTATACAAAGTTGG** |

**35) pGG-A-3xHA-B (from pEN-R2-3xHA-L3)**
**Colonies**: 1373 / **Efficiency**: 80%

| Vector | Position 1 | Position 2 |
|---|---|---|
| Original | TACAAAGTGGGTGGAGGCGGTTCAGCATGATACCCTTACGATGT | ACGACGTTCCAGATTACGCTTGATCAACTTTATTATACAAAGTTGG |
| Desired | AGAAGTGAAGCTTGGTCTCAACCTATGGCATACCCTTACGATGT | ACGACGTTCCAGATTACGCTTCAACATGAGACCGAATTCTCGCCCT |
| Clone 1 | AGAAGTGAAGCTTGGTCTCAACCTATGGCATACCCTTACGATGT | ACGACGTTCCAGATTAC-CTTCAACATGAGACCGAATTCTCGCCCT |
| Clone 2 | AGAAGTGAAGCTTGGTCTCAACCTATGGCATACCCTTACGATGT | ACGACGTTCCAGATTACGCTTCAACATGAGACCGAATTCTCGCCCT |
| Clone 3 | AGAAGTGAAGCTTGGTCTCAACCTATGGCATACCCTTACGATGT | ACGACGTTCCAGATTACGCTTCAACATGAGACCGAATTCTCGCCCT |
| Clone 4 | AGAAGTGAAGCTTGGTCTCAACCTATGGCATACCCTTACGATGT | ACGACGTTCCAGATTACGCTTCAACATGAGACCGAATTCTCGCCCT |
| Clone 5 | AGAAGTGAAGCTTGGTCTCAACCTATGGCATACCCTTACGATGT | ACGACGTTCCAGATTACGCTTCAACATGAGACCGAATTCTCGCCCT |

**36) pGG-B-3xHA-C (from pEN-R2-3xHA-L3)**
**Colonies**: 1584 / **Efficiency**: 80%

| Vector | Position 1 | Position 2 |
|---|---|---|
| Original | TACAAAGTGGGTGGAGGCGGTTCAGCATGATACCCTTACGATGT | ACGACGTTCCAGATTACGCTTGATCAACTTTATTATACAAAGTT |
| Desired | AGAAGTGAAGCTTGGTCTCAAACAATGGCATACCCTTACGATGT | ACGACGTTCCAGATTACGCTGGCTTGAGACCGAATTCTCGCCCT |
| Clone 1 | AGAAGTGAAGCTTGGTCTCAAACAATGGCATACCCTTACGATGT | ACGACGTTCCAGATTACGCTGGCTTGAGACCGAATTCTCGCCCT |
| Clone 2 | AGAAGTGAAGCTTGGTCTCAAACA-TGGCATACCCTTACGATGT | ACGACGTTCCAGATTACGCTGGCTTGAGACCGAATTCTCGCCCT |
| Clone 3 | AGAAGTGAAGCTTGGTCTCAAACAATGGCATACCCTTACGATGT | ACGACGTTCCAGATTACGCTGGCTTGAGACCGAATTCTCGCCCT |

### SUPPLEMENTARY MATERIALS

| | | |
|---|---|---|
| Clone 4 | AGAAGTGAAGCTTGGTCTCAAACAATGGCATACCCTTACGATGT | ACGACGTTCCAGATTACGCTGGCTTGAGACCGAATTCTCGCCCT |
| Clone 5 | AGAAGTGAAGCTTGGTCTCAAACAATGGCATACCCTTACGATGT | ACGACGTTCCAGATTACGCTGGCTTGAGACCGAATTCTCGCCCT |
| **37) pGG-C-3xHA-D (from pEN-R2-3xHA-L3)**<br>**Colonies**: 1845 / **Efficiency**: 100% | | |
| **Vector** | **Position 1** | **Position 2** |
| Original | TACAAAGTGGGTGGAGGCGGTTCAGCATGATACCCTTACGATGT | ACGACGTTCCAGATTACGCTTGATCAACTTTATTATACAAAGTT |
| Desired | AGAAGTGAAGCTTGGTCTCAGGCTCTATGGCATACCCTTACGATGT | ACGACGTTCCAGATTACGCTTCAGTGAGACCGAATTCTCGCCCT |
| Clone 1 | AGAAGTGAAGCTTGGTCTCAGGCTCTATGGCATACCCTTACGATGT | ACGACGTTCCAGATTACGCTTCAGTGAGACCGAATTCTCGCCCT |
| Clone 2 | AGAAGTGAAGCTTGGTCTCAGGCTCTATGGCATACCCTTACGATGT | ACGACGTTCCAGATTACGCTTCAGTGAGACCGAATTCTCGCCCT |
| Clone 3 | AGAAGTGAAGCTTGGTCTCAGGCTCTATGGCATACCCTTACGATGT | ACGACGTTCCAGATTACGCTTCAGTGAGACCGAATTCTCGCCCT |
| Clone 4 | AGAAGTGAAGCTTGGTCTCAGGCTCTATGGCATACCCTTACGATGT | ACGACGTTCCAGATTACGCTTCAGTGAGACCGAATTCTCGCCCT |
| Clone 5 | AGAAGTGAAGCTTGGTCTCAGGCTCTATGGCATACCCTTACGATGT | ACGACGTTCCAGATTACGCTTCAGTGAGACCGAATTCTCGCCCT |
| **38) pGG-D-3xHA-E (from pEN-R2-3xHA-L3)**<br>**Colonies**: 936 / **Efficiency**: 80% | | |
| **Vector** | **Position 1** | **Position 2** |
| Original | TACAAAGTGGGTGGAGGCGGTTCAGCATGATACCCTTACGATGT | ACGACGTTCCAGATTACGCTTGATCAACTTTATTATACAAAGTT |
| Desired | AGAAGTGAAGCTTGGTCTCATCAGCTATGGCATACCCTTACGATGT | ACGACGTTCCAGATTACGCTCTGCTGAGACCGAATTCTCGCCCT |
| Clone 1 | AGAAGTGAAGCTTGGTCTCATCAGCTATGGCATACCCTTACGATGT | ACGACGTTCCAGATTACGCTCTGCTGAGACCGAATTCTCGCCCT |
| Clone 2 | AGAAGTGAAGCTTGGTCTCATCAGCTATGGCATACCCTTACGATGT | ACGACGTTCCAGATTACGCTCTGCTGAGACCGAATTCTCGCCCT |
| Clone 3 | AGAAGTGAAGCTTGGTCTCATCAGCTATGGCATACCCTTACGATGT | ACGACGTTCCAGATTACGCTCTGCTGAGACCGAATTCTCGCCCT |
| Clone 4 | AGAAGTGAAGCTTGGTCTCATCAGCTATGGCATACCCTTACGATGT | ACGACGTTCCAGATTACGCTCTGCTGAGACCGAATTCTCGCCCT |
| Clone 5 | AGAAGTGAAGCTTGGTCTCATCAGCTATGGCATACCCTTACGATGT † | ACGACGTTCCAGATTACGCTCTGCTGAGACCGAATTCTCGCCCT |
| **39) pGG-E-3xHA-F (from pEN-R2-3xHA-L3)**<br>**Colonies**: 1257 / **Efficiency**: 100% | | |
| **Vector** | **Position 1** | **Position 2** |
| Original | TACAAAGTGGGTGGAGGCGGTTCAGCATGATACCCTTACGATGT | ACGACGTTCCAGATTACGCTTGATCAACTTTATTATACAAAGTTGGC |
| Desired | AGAAGTGAAGCTTGGTCTCACTGCCTATGGCATACCCTTACGATGT | ACGACGTTCCAGATTACGCTTGAACTATGAGACCGAATTCTCGCCCT |
| Clone 1 | AGAAGTGAAGCTTGGTCTCACTGCCTATGGCATACCCTTACGATGT | ACGACGTTCCAGATTACGCTTGAACTATGAGACCGAATTCTCGCCCT |
| Clone 2 | AGAAGTGAAGCTTGGTCTCACTGCCTATGGCATACCCTTACGATGT | ACGACGTTCCAGATTACGCTTGAACTATGAGACCGAATTCTCGCCCT |
| Clone 3 | AGAAGTGAAGCTTGGTCTCACTGCCTATGGCATACCCTTACGATGT | ACGACGTTCCAGATTACGCTTGAACTATGAGACCGAATTCTCGCCCT |
| Clone 4 | AGAAGTGAAGCTTGGTCTCACTGCCTATGGCATACCCTTACGATGT | ACGACGTTCCAGATTACGCTTGAACTATGAGACCGAATTCTCGCCCT |
| Clone 5 | AGAAGTGAAGCTTGGTCTCACTGCCTATGGCATACCCTTACGATGT | ACGACGTTCCAGATTACGCTTGAACTATGAGACCGAATTCTCGCCCT |

*attL3

† part of 3xHA is missing

**SUPPLEMENTARY MATERIALS**

### Supplementary Table 6: Clone tagging with oligo-encoded sequences

Partial sequences of the original sequence, the desired sequence and five sequenced clones are shown in each table. The positions in the table refer to the junctions in final tagged clone, for the original untagged vector the end that is tagged is shown twice in the table. The BsaI recognition motif is underlined. The 4-bp overhangs that are generated by BsaI digestion are shown in bold. The sequence that is tagged is highlighted in blue, the tag in green. The sequence highlighted in gray is the backbone sequence. Mutations that deviate from the desired sequence are highlighted in red. The number of colonies and the cloning efficiency is also indicated. Experiment number is indicated in top left corner.

| **40) ER signal peptide-Integrase 4 fusion (oligo-encoded) to pGGC0000 (GmR)**<br>**Colonies:** 118 / **Efficiency**: 80% | |
|---|---|
| **Vector** | **Position 1** |
| Original | AGAAGTGAAGCTTGGTCTCA**GGCT**CC ATGATTACGACCAGAAAG |
| Desired | AGAAGTGAAGCTTGGTCTCA**GGCT**CCATGAAGGTACAGGAGGGTTTGTTCGTGGTGGCTGTTTTCTACCTTGCTTATACGCAGCTAGTCAAGGGGATGATTACGACCAGAAAG |
| Clone 1 | AGAAGTGAAGCTTGGTCTCA**GGCT**CCATGAAGGTACAGGAGGGTTTGTTCGTGGTGGCTGTTTTCTACCTTGCTTATACGCAGCTAGTCAAGGGGATGATTACGACCAGAAAG |
| Clone 2 | AGAAGTGAAGCTTGGTCTCA**GGCT**CCATGAAGGTACAGGAGGGTTTGTTCGTGGTGGCTGTTTTCTACCTTGCTTATACGCAGCTAGTCAAGGGGATGATTACGACCAGAAAG |
| Clone 3 | AGAAGTGAAGCTTGGTCTCA**GGCT**CCATGAAGGTACAGGAGGGTTTGTTCGTGGTGGCTGTTTTCTACCTTGCTTATACGCAGCTAGTCAAGGGGATGATTACGACCAGAAAG |
| Clone 4 | AGAAGTGAAGCTTGGTCTCA**GGCT**CCATGAAGGTACAGGAGGGTTTGTTCGTGGTGGCTGTTTTCTACCTTGCTTATACGCAGCTAGTCAAGGGGATGATTACGACCAGAAAG |
| Clone 5 | AGAAGTGAAGCTTGGTCT---GCTCCATGAAGGTACAGGAGGGTTTGTTCGTGGTGGCTGTTTTCTACCTTGCTTATACGCAGCTAGTCAAGGGGATGATTACGACCAGAAAG |
| **Vector** | **Position 2** |
| Original | TAGATATACATTGGACCTTT**TCAG**TGAGACCGAATTCTCGCCCT |
| Desired | TAGATATACATTGGACCTTT**TCAG**TGAGACCGAATTCTCGCCCT |
| Clone 1 | TAGATATACATTGGACCTTT**TCAG**TGAGACCGAATTCTCGCCCT |
| Clone 2 | TAGATATACATTGGACCTTT**TCAG**TGAGACCGAATTCTCGCCCT |
| Clone 3 | TAGATATACATTGGACCTTT**TCAG**TGAGACCGAATTCTCGCCCT |
| Clone 4 | TAGATATACATTGGACCTTT**TCAG**TGAGACCGAATTCTCGCCCT |
| Clone 5 | TAGATATACATTGGACCTTT**TCAG**TGAGACCGAATTCTCGCCCT |

| **41) Integrase 4-SV40 fusion (oligo-encoded) to pGGC0000 (TetR)**<br>**Colonies:** 29 / **Efficiency**: 100% | |
|---|---|
| **Vector** | **Position 1** |
| Original | AGAAGTGAAGCTTGGTCTCA**GGCT**CCATGATTACGACCAGAAAG |
| Desired | AGAAGTGAAGCTTGGTCTCA**GGCT**CCATGATTACGACCAGAAAG |
| Clone 1 | AGAAGTGAAGCTTGGTCTCA**GGCT**CCATGATTACGACCAGAAAG |
| Clone 2 | AGAAGTGAAGCTTGGTCTCA**GGCT**CCATGATTACGACCAGAAAG |
| Clone 3 | AGAAGTGAAGCTTGGTCTCA**GGCT**CCATGATTACGACCAGAAAG |
| Clone 4 | AGAAGTGAAGCTTGGTCTCA**GGCT**CCATGATTACGACCAGAAAG |
| Clone 5 | AGAAGTGAAGCTTGGTCTCA**GGCT**CCATGATTACGACCAGAAAG |
| **Vector** | **Position 2** |
| Original | TAGATATACATTGGACCTTT **TCAG**TGAGACCGAATTCTCGCCCT |

| | |
|---|---|
| Desired | TAGATATACATTGGACCTTTCCTAAGAAGAAGAGGAAGGTT**TCAG**TGAGACCGAATTCTCGCCCT |
| Clone 1 | TAGATATACATTGGACCTTTCCTAAGAAGAAGAGGAAGGTT**TCAG**TGAGACCGAATTCTCGCCCT |
| Clone 2 | TAGATATACATTGGACCTTTCCTAAGAAGAAGAGGAAGGTT**TCAG**TGAGACCGAATTCTCGCCCT |
| Clone 3 | TAGATATACATTGGACCTTTCCTAAGAAGAAGAGGAAGGTT**TCAG**TGAGACCGAATTCTCGCCCT |
| Clone 4 | TAGATATACATTGGACCTTTCCTAAGAAGAAGAGGAAGGTT**TCAG**TGAGACCGAATTCTCGCCCT |
| Clone 5 | TAGATATACATTGGACCTTTCCTAAGAAGAAGAGGAAGGTT**TCAG**TGAGACCGAATTCTCGCCCT |

### 42) ER signal peptide-Integrase 4-SV40 fusion (oligo-encoded) to pGGC0000 (SpecR)

**Colonies:** 45 / **Efficiency**: 80%

| **Vector** | **Position 1** |
|---|---|
| Original | AGAAGTGAAGCTTGGTCTCA**GGCT**CC ATGATTACGACCAGAAAG |
| Desired | AGAAGTGAAGCTTGGTCTCA**GGCT**CCATGAAGGTACAGGAGGGTTTGTTCGTGGTGGCTGTTTTCTACCTTGCTTATACGCAGCTAGTCAAGGGGATGATTACGACCAGAAAG |
| Clone 1 | AGAAGTGAAGCTTGGTCTCA**GGCT**CCATGAAGGTACAGGAGGGTTTGTTCGTGGTGGCTGTTTTCTACCTTGCTTATACGCAGCTAGTCAAGGGGATGATTACGACCAGAAAG |
| Clone 2 | AGAAGTGAAGCTTGGTCTCA**GGCT**CCATGAAGGTACAGGAGGGTTTGTTCGTGGTGGCTGTTTTCTACCTTGCTTATACGCAGCTAGTCAAGGGGATGATTACGACCAGAAAG |
| Clone 3 | AGAAGTGAAGCTTGGTCTCA**GGCT**CCATGAAGGTACAGGAGGGTTTGTTCGTGGTGGCTGTTTTCTACCTTGCTTATAC-CAGCTAGTCAAGGGGATGATTACGACCAGAAAG |
| Clone 4 | AGAAGTGAAGCTTGGTCTCA**GGCT**CCATGAAGGTACAGGAGGGTTTGTTCGTGGTGGCTGTTTTCTACCTTGCTTATACGCAGCTAGTCAAGGGGATGATTACGACCAGAAAG |
| Clone 5 | AGAAGTGAAGCTTGGTCTCA**GGCT**CCATGAAGGTACAGGAGGGTTTGTTCGTGGTGGCTGTTTTCTACCTTGCTTATACGCAGCTAGTCAAGGGGATGATTACGACCAGAAAG |
| **Vector** | **Position 2** |
| Original | TAGATATACATTGGACCTTT **TCAG**TGAGACCGAATTCTCGCCCT |
| Desired | TAGATATACATTGGACCTTTCCTAAGAAGAAGAGGAAGGTT**TCAG**TGAGACCGAATTCTCGCCCT |
| Clone 1 | TAGATATACATTGGACCTTTCCTAAGAAGAAGAGGAAGGTT**TCAG**TGAGACCGAATTCTCGCCCT |
| Clone 2 | TAGATATACATTGGACCTTTCCTAAGAAGAAGAGGAAGGTT**TCAG**TGAGACCGAATTCTCGCCCT |
| Clone 3 | TAGATATACATTGGACCTTTCCTAAGAAGAAGAGGAAGGTT**TCAG**TGAGACCGAATTCTCGCCCT |
| Clone 4 | TAGATATACATTGGACCTTTCCTAAGAAGAAGAGGAAGGTT**TCAG**TGAGACCGAATTCTCGCCCT |
| Clone 5 | TAGATATACATTGGACCTTTCCTAAGAAGAAGAGGAAGGTT**TCAG**TGAGACCGAATTCTCGCCCT |

**SUPPLEMENTARY MATERIALS**

Supplementary Table 7: Restriction enzyme recognition site removal

Partial sequences of the original sequence, the desired sequence and five sequenced clones are shown in each table. The restriction enzyme recognition sites are shown in bold, flanked by backbone at both sides. The base that is targeted for replacement is highlighted in blue in the original sequence and in green in the desired final sequence. Mutations that deviate from the desired sequence are highlighted in red. The number of colonies and the cloning efficiency is also indicated. Experiment number is indicated in top left corner.

**47) Removal of XbaI in A-OLE-P-B**
**Colonies:** 673 / **Efficiency**: 100%

| Vector | Position 1 (2525-2568) | Position 2 (3162-3205) |
|---|---|---|
| Original | CGATCTGATACTGATAACG**TCTAGA**TTTTTAGGGTTAAAGCAAT | TATCCATTTTCTTCATTGT**TCTAGA**ATGTCGCGGAACAAATTTT |
| Desired | CGATCTGATACTGATAACG**TCTCGA**TTTTTAGGGTTAAAGCAAT | TATCCATTTTCTTCATTGT**TATAGA**ATGTCGCGGAACAAATTTT |
| Clone 1 | CGATCTGATACTGATAACG**TCTCGA**TTTTTAGGGTTAAAGCAAT | TATCCATTTTCTTCATTGT**TATAGA**ATGTCGCGGAACAAATTTT |
| Clone 2 | CGATCTGATACTGATAACG**TCTCGA**TTTTTAGGGTTAAAGCAAT | TATCCATTTTCTTCATTGT**TATAGA**ATGTCGCGGAACAAATTTT |

**48) Removal of EcorI in B-Csy4-C**
**Colonies:** 97 / **Efficiency**: 60%

| Vector | Position 1 (199-242) | Position 2 (412-455) | Position 3 (662-705) |
|---|---|---|---|
| Original | AGACTTGGAGAAAGACTTA**GAATTC**ATGCTTCTGCTGATGATCT | GAAGAAGAAGCTAGAAAAA**GAATTC**CTGATACTGTTGCTAGAGC | CTGGACCCGGCTTGAGACC**GAATTC**TCGCCCTATAGTGAGTCGT |
| Desired | AGACTTGGAGAAAGACTTA**GGATTC**ATGCTTCTGCTGATGATCT | GAAGAAGAAGCTAGAAAAA**GAGTTC**CTGATACTGTTGCTAGAGC | CTGGACCCGGCTTGAGACC**GAACTC**TCGCCCTATAGTGAGTCGT |
| Clone 1 | AGACTTGGAGAAAGACTTA**G-ATTC**ATGCTTCTGCTGATGATCT | GAAGAAGAAGCTAGAAAAA**GAGTTC**CTGATACTGTTGCTAGAGC | CTGGACCCGGCTTGAGACC**GAACTC**TCGCCCTATAGTGAGTCGT |
| Clone 2 | AGACTTGGAGAAAGACTTA**GGATTC**ATGCTTCTGCTGATGATCT | GAAGAAGAAGCTAGAAAAA**GAGTTC**CTGATACTGTTGCTAGAGC | CTGGACCCGGCTTGAGACC**GAACTC**TCGCCCTATAGTGAGTCGT |
| Clone 3 | AGACTTGGAGAAAGACTTA**GGATTC**ATGCTTCTGCTGATGATCT | GAAGAAGAAGCTAGAAAAA**GAGTTC**CTGATACTGTTGCTAGAGC | CTGGACCCGGCTTGAGACC**GAACTC**TCGCCCTATAGTGAGTCGT |
| Clone 4 | AGACTTGGAGAAAGACTTA**GGATTC**ATGCTTCTGCTGATGATCT | GAAGAAGAAGCTAGAAAAA**GAGTTC**CTGATACTGTTGCTAGAGC | CTGGACCCGGCTTGAGACC**GAACTC**TCGCCCTATAGTGAGTCGT |
| Clone 5 | AGACTTGGAGAAAGACTTA**GGATTC**ATGCTTCTGCTGATGATCT | AAGAAGAAGCTAGGAAAAA**GAGTTC**CTGATACTGTTGCTAGAGC | CTGGACCCGGCTTGAGACC**GAACTC**TCGCCCTATAGTGAGTCGT |

**49) Remove extra BsaI site from pGG-E-tUM140_0016-F*** (Oligo 2a)**
**Colonies**: 189 / **Efficiency**: 0%

| Vector | Position 1 | Position 2 | Position 3 |
|---|---|---|---|
| Original | AGAAGTGAAGCTTGGTCTCACTGCGCCGCGCACAGCTGACGTAG | AACTCTAGTGTAATCCCAGACTGGTGAGACCAGTGACATTGACACCA | TACATTTGCATGCAAAGTTGACTATGAGACCGAATTCTCGCCCT |
| Desired | AGAAGTGAAGCTTGGTCTCACTGCGCCGCGCACAGCTGACGTAG | AACTCTAGTGTAATCCCAGACTGGTGTGACCAGTGACATTGACACCA | TACATTTGCATGCAAAGTTGACTATGAGACCGAATTCTCGCCCT |
| Clone 1 | AGAAGTGAAGCTTGGTCTCACTGCGCCGCGCACAGCTGACGTAG | AACTCTAGTGTAATCCCAGACTGGTGAGACCAGTGACATTGACACCA | TACATTTGCATGCAAAGTTGACTATGAGACCGAATTCTCGCCCT |
| Clone 2 | AGAAGTGAAGCTTGGTCTCACTGCGCCGCGCACAGCTGACGTAG | AACTCTAGTGTAATCCCAGACTGGTGAGACCAGTGACATTGACACCA | TACATTTGCATGCAAAGTTGACTATGAGACCGAATTCTCGCCCT |
| Clone 3 | AGAAGTGAAGCTTGGTCTCACTGCGCCGCGCACAGCTGACGTAG | AACTCTAGTGTAATCCCAGACTGGTGAGACCAGTGACATTGACACCA | TACATTTGCATGCAAAGTTGACTATGAGACCGAATTCTCGCCCT |
| Clone 4 | AGAAGTGAAGCTTGGTCTCACTGCGCCGCGCACAGCTGACGTAG | AACTCTAGTGTAATCCCAGACTGGTGAGACCAGTGACATTGACACCA | TACATTTGCATGCAAAGTTGACTATGAGACCGAATTCTCGCCCT |
| Clone 5 | AGAAGTGAAGCTTGGTCTCACTGCGCCGCGCACAGCTGACGTAG | AACTCTAGTGTAATCCCAGACTGGTGAGACCAGTGACATTGACACCA | TACATTTGCATGCAAAGTTGACTATGAGACCGAATTCTCGCCCT |

**50) Remove extra BsaI site from pGG-E-tUM140_0016-F (Oligo 2b + GmR)**
**Colonies**: 11 / **Efficiency**: 40%

| Vector | Left Flank | Extra BsaI site | Right Flank |
|---|---|---|---|

| | | | |
|---|---|---|---|
| Original | AGAAGTGAAGCTTGGTCTCACTGCGCCGCGCACAGCTGACGTAG | AACTCTAGTGTAATCCCAGACTGGTGAGACCAGTGACATTGACACCA | TACATTTGCATGCAAAGTTGACTATGAGACCGAATTCTCGCCCT |
| Desired | AGAAGTGAAGCTTGGTCTCACTGCGCCGCGCACAGCTGACGTAG | AACTCTAGTGTAATCCCAGACTGGTGAGGCCAGTGACATTGACACCA | TACATTTGCATGCAAAGTTGACTATGAGACCGAATTCTCGCCCT |
| Clone 1 | Backbone | Backbone | Backbone |
| Clone 2 | Backbone | Backbone | Backbone |
| Clone 3 | AGAAGTGAAGCTTGGTCTCACTGCGCCGCGCACAGCTGACGTAG | AACTCTAGTGTAATCCCAGACTGGTGAGGCCAGTGACATTGACACCA | TACATTTGCATGCAAAGTTGACTATGAGACCGAATTCTCGCCCT |
| Clone 4 | AGAAGTGAAGCTTGGTCTCACTGCGCCGCGCACAGCTGACGTAG | AACTCTAGTGTAATCCCAGACTGGTGAGACCAGTGACATTGACACCA | TACATTTGCATGCAAAGTTGACTATGAGACCGAATTCTCGCCCT |
| Clone 5 | AGAAGTGAAGCTTGGTCTCACTGCGCCGCGCACAGCTGACGTAG | AACTCTAGTGTAATCCCAGACTGGTGAGGCCAGTGACATTGACACCA | TACATTTGCATGCAAAGTTGACTATGAGACCGAATTCTCGCCCT |

**51) Remove extra BsaI site from pGG-E-tUM140_0016-F (Oligo 2d + GmR)**

**Colonies**: 12 / **Efficiency**: 60%

| Vector | Left Flank | Extra BsaI site | Right Flank |
|---|---|---|---|
| Original | AGAAGTGAAGCTTGGTCTCACTGCGCCGCGCACAGCTGACGTAG | AACTCTAGTGTAATCCCAGACTGGTGAGACCAGTGACATTGACACCA | TACATTTGCATGCAAAGTTGACTATGAGACCGAATTCTCGCCCT |
| Desired | AGAAGTGAAGCTTGGTCTCACTGCGCCGCGCACAGCTGACGTAG | AACTCTAGTGTAATCCCAGACTGGTGAGAACAGTGACATTGACACCA | TACATTTGCATGCAAAGTTGACTATGAGACCGAATTCTCGCCCT |
| Clone 1 | AGAAGTGAAGCTTGGTCTCACTGCGCCGCGCACAGCTGACGTAG | AACTCTAGTGTAATCCCAGACTGGTGAGAACAGTGACATTGACACCA | TACATTTGCATGCAAAGTTGACTATGAGACCGAATTCTCGCCCT |
| Clone 2 | AGAAGTGAAGCTTGGTCTCACTGCGCCGCGCACAGCTGACGTAG | AACTCTAGTGTAATCCCAGACTGGTGAGAACAGTGACATTGACACCA | TACATTTGCATGCAAAGTTGACTATGAGACCGAATTCTCGCCCT |
| Clone 3 | AGAAGTGAAGCTTGGTCTCACTGCGCCGCGCACAGCTGACGTAG | AACTCTAGTGTAATCCCAGACTGGTGAGAACAGTGACATTGACACCA | TACATTTGCATGCAAAGTTGACTATGAGACCGAATTCTCGCCCT |
| Clone 4 | AGAAGTGAAGCTTGGTCTCACTGCGCCGCGCACAGCTGACGTAG | AACTCTAGTGTAATCCCAGACTGGTGAGACCAGTGACATTGACACCA | TACATTTGCATGCAAAGTTGACTATGAGACCGAATTCTCGCCCT |
| Clone 5 | Backbone | Backbone | Backbone |

**52) mCherry: remove BbsI + transfer to Moclo (pGGC0000 SpecR)**

**Colonies**: 496 / **Efficiency**: 40%

| Vector | Left Flank | BbsI site | Right Flank |
|---|---|---|---|
| Original | AGAAGTGAAGCTTGGTCTCATCAGGTATGGTGAGCAAGGGCGAGGA | GGCCCCGTAATGCAGAAGAAGACCATGGGCTGGGAGGCCTCCTCCGAGC | GCATGGACGAGCTGTACAAGCTGCTGAGACCGAATTCTCGCCCT |
| Desired | AGAAGTGAAGCTTGGTCTCATTCGCTATGGTGAGCAAGGGCGAGGA | GGCCCCGTAATGCAGAAGAAAACCATGGGCTGGGAGGCCTCCTCCGAGC | GCATGGACGAGCTGTACAAGGCTTTGAGACCGAATTCTCGCCCT |
| Clone 1 | AGAAGTGAAGCTTGGTCTCATTCGCTATGGTGAGCAAGGGCGAGGA | GGCCCCGTAATGCAGAAGAAAACCATGGGCTGGGAGGCCTCCTCCGAGC | GCATGGACGAGCTGTACAAGGCTTTGAGACCGAATTCTCGCCCT |
| Clone 2 | AGAAGTGAAGCTTGGTCTCATTCGCTATGGTGAGCAAGGGCGAGGA | GGCCCCGTAATGCAGAAGAAAACCATGGGCTGGGAGGCCTCCTCCGAGC | GCATGGACGAGCTGTACAAGGCTTTGAGACCGAATTCTCGCCCT |
| Clone 3 | AGAAGTGAAGCTTGGTCTCATTCGCTATGGTGAGCAAGGGCGAGGA | GGCCCCGTAATGCAGAAGAAGACCATGGGCTGGGAGGCCTCCTCCGAGC | Backbone |
| Clone 4 | AGAAGTGAAGCTTGGTCTCATTCGCTATGGTGAGCAAGGGCGAGGA | GGCCCCGTAATGCAGAAGAAGACCATGGGCTGGGAGGCCTCCTCCGAGC | GCATGGACGAGCTGTACAAGGCTTTGAGACCGAATTCTCGCCCT |
| Clone 5 | AGAAGTGAAGCTTGGTCTCATTCGCTATGGTGAGCAAGGGCGAGGA | GGCCCCGTAATGCAGAAGAAGACCATGGGCTGGGAGGCCTCCTCCGAGC | Backbone |

**53) WUS: remove BbsI + transfer to Moclo pICH41233 (blue-white screen)**

**Colonies**: 24 white, 19 blue/ **Efficiency**: 20%

| Vector | Left Flank | BbsI site | Right Flank |
|---|---|---|---|
| Original | AGAAGTGAAGCTTGGTCTCAACCTCCCATGTTTACGTTTACGTT | TACTCAACATGTTCATAAGTACACCTGTCTTCACACTCGTTTCACACA | CAAAAGTCGAATCAAACACACAACATGAGACCGAATTCTCGCCCT |
| Desired | TTAATCACTCTGTGGTCTCAGGAGCCCATGTTTACGTTTACGTT | TACTCAACATGTTCATAAGTACACCTGACTTCACACTCGTTTCACACA | CAAAAGTCGAATCAAACACACCCATTGAGACCACGAAGTGGCTCT |
| Clone 1 | TTAATCACTCTGTGGTCTCAGGAGCCCATGTTTACGTTTACGTT | TACTCAACATGTTCATAAGTACACCTGTCTTCACACTCGTTTCACACA | CAAAAGTCGAATCAAACACACCCATTGAGACCACGAAGTGGCTCT |
| Clone 2 | TTAATCACTCTGTGGTCTCAGGAGCCCATGTTTACGTTTACGTT | TACTCAACATGTTCATAAGTACACCTGACTTCACACTCGTTTCACACA | CAAAAGTCGAATCAAACACACCCATTGAGACCACGAAGTGGCTCT |
| Clone 3 | TTAATCACTCTGTGGTCTCAGGAGCCCATGTTTACGTTTACGTT | TACTCAACATGTTCATAAGTACACCTGTCTTCACACTCGTTTCACACA | CAAAAGTCGAATCAAACACACCCATTGAGACCACGAAGTGGCTCT |
| Clone 4 | TTAATCACTCTGTGGTCTCAGGAGCCCATGTTTACGTTTACGTT | TACTCAACATGTTCATAAGTACACCTGTCTTCACACTCGTTTCACACA | CAAAAGTCGAATCAAACACACCCATTGAGACCACGAAGTGGCTCT |
| Clone 5 | TTAATCACTCTGTGGTCTCAGGAGCCCATGTTTACGTTTACGTT | TACTCAACATGTTCATAAGTACACCTGTCTTCACACTCGTTTCACACA | CAAAAGTCGAATCAAACACACCCATTGAGACCACGAAGTGGCTCT |

**54) WUS: remove BbsI + transfer to Moclo (with SpecR alternative)**

### SUPPLEMENTARY MATERIALS

**Colonies**: 548 / **Efficiency**: 0%

| Vector | Left Flank | BbsI site | Right Flank |
|---|---|---|---|
| Original | AGAAGTGAAGCTTGGTCTCAACCTCCCATGTTTACGTTTACGTT | TACTCAACATGTTCATAAGTACACCTGTCTTCACACTCGTTTCACACA | CAAAAGTCGAATCAAACACACAACATGAGACCGAATTCTCGCCCT |
| Desired | AGAAGTGAAGCTTGGTCTCAGGAGCCCATGTTTACGTTTACGTT | TACTCAACATGTTCATAAGTACACCTGACTTCACACTCGTTTCACACA | CAAAAGTCGAATCAAACACACCCATTGAGACCGAATTCTCGCCCT |
| Clone 1 | AGAAGTGAAGCTTGGTCTCAGGAGCCCATGTTTACGTTTACGTT | TACTCAACATGTTCATAAGTACACCTGTCTTCACACTCGTTTCACACA | CAAAAGTCGAATCAAACACACCCATTGAGACCGAATTCTCGCCCT |
| Clone 2 | AGAAGTGAAGCTTGGTCTCAGGAGCCCATGTTTACGTTTACGTT | TACTCAACATGTTCATAAGTACACCTGTCTTCACACTCGTTTCACACA | CAAAAGTCGAATCAAACACACCCATTGAGACCGAATTCTCGCCCT |
| Clone 3 | AGAAGTGAAGCTTGGTCTCAGGAGCCCATGTTTACGTTTACGTT | TACTCAACATGTTCATAAGTACACCTGTCTTCACACTCGTTTCACACA | CAAAAGTCGAATCAAACACACCCATTGAGACCGAATTCTCGCCCT |
| Clone 4 | AGAAGTGAAGCTTGGTCTCAGGAGCCCATGTTTACGTTTACGTT | TACTCAACATGTTCATAAGTACACCTGTCTTCACACTCGTTTCACACA | CAAAAGTCGAATCAAACACACCCATTGAGACCGAATTCTCGCCCT |
| Clone 5 | AGAAGTGAAGCTTGGTCTCAGGAGCCCATGTTTACGTTTACGTT | TACTCAACATGTTCATAAGTACACCTGTCTTCACACTCGTTTCACACA | CAAAAGTCGAATCAAACACACCCATTGAGACCGAATTCTCGCCCT |

#### 55) pGG-D-GUS-E: remove both AarI sites

**Colonies**: 38 / **Efficiency**: 10%

| Vector | Position 1 | Position 2 |
|---|---|---|
| Original | AAGACTGTAACCACGCGTCTGTTGACTGGCAGGTGGTGGCCAATGGTGATGT | CCGACGCGTCCGATCACCTGCGTCAATGTAATGTTCTGCGACGCTCACA |
| Desired | AAGACTGTAACCACGCGTCTGTTGACTGGCAAGTGGTGGCCAATGGTGATGT | CCGACGCGTCCGATCACCTGTGTCAATGTAATGTTCTGCGACGCTCACA |
| Clone 1 | AAGACTGTAACCACGCGTCTGTTGACTGGCAGGTGGTGGCCAATGGTGATGT | CCGACGCGTCCGATCACCTGTGTCAATGTAATGTTCTGCGACGCTCACA |
| Clone 2 | AAGACTGTAACCACGCGTCTGTTGACTGGCAGGTGGTGGCCAATGGTGATGT | CCGACGCGTCCGATCACCTGTGTCAATGTAATGTTCTGCGACGCTCACA |
| Clone 3 | AAGACTGTAACCACGCGTCTGTTGACTGGCAAGTGGTGGCCAATGGTGATGT | CCGACGCGTCCGATCACCTGTGTCAATGTAATGTTCTGCGACGCTCACA |
| Clone 4 | AAGACTGTAACCACGCGTCTGTTGACTGGCAGGTGGTGGCCAATGGTGATGT | CCGACGCGTCCGATCACCTGTGTCAATGTAATGTTCTGCGACGCTCACA |
| Clone 5 | AAGACTGTAACCACGCGTCTGTTGACTGGCAGGTGGTGGCCAATGGTGATGT | CCGACGCGTCCGATCACCTGCGTCAATGTAATGTTCTGCGACGCTCACA |
| Clone 6 | AAGACTGTAACCACGCGTCTGTTGACTGGCAGGTGGTGGCCAATGGTGATGT | CCGACGCGTCCGATCACCTGTGTCAATGTAATGTTCTGCGACGCTCACA |
| Clone 7 | AAGACTGTAACCACGCGTCTGTTGACTGGCAGGTGGTGGCCAATGGTGATGT | CCGACGCGTCCGATCACCTGTGTCAATGTAATGTTCTGCGACGCTCACA |
| Clone 8 | AAGACTGTAACCACGCGTCTGTTGACTGGCAGGTGGTGGCCAATGGTGATGT | CCGACGCGTCCGATCACCTGTGTCAATGTAATGTTCTGCGACGCTCACA |
| Clone 9 | AAGACTGTAACCACGCGTCTGTTGACTGGCAGGTGGTGGCCAATGGTGATGT | CCGACGCGTCCGATCACCTGTGTCAATGTAATGTTCTGCGACGCTCACA |
| Clone 10 | AAGACTGTAACCACGCGTCTGTTGACTGGCAGGTGGTGGCCAATGGTGATGT | CCGACGCGTCCGATCACCTGTGTCAATGTAATGTTCTGCGACGCTCACA |

#### 56) pGG-D-GUS-E: remove both AarI sites (Oligo Aar1b)

**Colonies**: 20 / **Efficiency**: 20%

| Vector | Position 1 | Position 2 |
|---|---|---|
| Original | TGTAACCACGCGTCTGTTGACTGGCAGGTGGTGGCCAATGGTGATGTCAG | CCGACGCGTCCGATCACCTGCGTCAATGTAATGTTCTGCGACGC |
| Desired | TGTAACCACGCGTCTGTTGACTGGCAGGTCGTGGCCAATGGTGATGTCAG | CCGACGCGTCCGATCACCTGTGTCAATGTAATGTTCTGCGACGC |
| Clone 1 | TGTAACCACGCGTCTGTTGACTGGCAGGTCGTGGCCAATGGTGATGTCAG | CCGACGCGTCCGATCACCTGTGTCAATGTAATGTTCTGCGACGC |
| Clone 2 | TGTAACCACGCGTCTGTTGACTGGCAGGTGGTGGCCAATGGTGATGTCAG | CCGACGCGTCCGATCACCTGTGTCAATGTAATGTTCTGCGACGC |
| Clone 3 | TGTAACCACGCGTCTGTTGACTGGCAGGTGGTGGCCAATGGTGATGTCAG | CCGACGCGTCCGATCACCTGTGTCAATGTAATGTTCTGCGACGC |
| Clone 4 | TGTAACCACGCGTCTGTTGACTGGCAGGTGGTGGCCAATGGTGATGTCAG | CCGACGCGTCCGATCACCTGTGTCAATGTAATGTTCTGCGACGC |
| Clone 5 | TGTAACCACGCGTCTGTTGACTGGCAGGTGGTGGCCAATGGTGATGTCAG | CCGACGCGTCCGATCACCTGTGTCAATGTAATGTTCTGCGACGC |

**SUPPLEMENTARY MATERIALS**

Supplementary Table 8: Genotype and color phenotype of selected colonies

Table showing the partial vector sequence (i.e., the six bases that were mutagenized flanked by 20 bp on each end) of the original vector, compared with the 55 picked clones of Supplementary Figure 4.

| Vector | Sequence | AA identity (64-65) | Color |
|---|---|---|---|
| Original | GGGATATTTTATCACCACAG**GTTGGA**TACGGAAGCATACCATTCAC | VG | |
| Clone 1 | GGGATATTTTATCACCACAG**TTTTTA**TACGGAAGCATACCATTCAC | FL | |
| Clone 2 | GGGATATTTTATCACCACAG**CTATGT**TACGGAAGCATACCATTCAC | LC | |
| Clone 3 | GGGATATTTTATCACCACAG**ATGTCT**TACGGAAGCATACCATTCAC | MS | |
| Clone 4 | GGGATATTTTATCACCACAG**TTATGC**TACGGAAGCATACCATTCAC | LC | |
| Clone 5 | GGGATATTTTATCACCACAG**TATACG**TACGGAAGCATACCATTCAC | YT | |
| Clone 6 | GGGATATTTTATCACCACAG**ATTGCC**TACGGAAGCATACCATTCAC | IA | |
| Clone 7 | GGGATATTTTATCACCACAG**TTTGTA**TACGGAAGCATACCATTCAC | FV | |
| Clone 8 | GGGATATTTTATCACCACAG**AATTCT**TACGGAAGCATACCATTCAC | NS | |
| Clone 9 | GGGATATTTTATCACCACAG**TGGTGT**TACGGAAGCATACCATTCAC | WC | |
| Clone 10 | GGGATATTTTATCACCACAG**TGGACC**TACGGAAGCATACCATTCAC | WT | |
| Clone 11 | GGGATATTTTATCACCACAG**TTAATG**TACGGAAGCATACCATTCAC | LM | |
| Clone 12 | GGGATATTTTATCACCACAG**TTAATG**TACGGAAGCATACCATTCAC | LM | |
| Clone 13 | GGGATATTTTATCACCACAG**CTGATG**TACGGAAGCATACCATTCAC | LM | |
| Clone 14 | GGGATATTTTATCACCACAG**TTCATG**TACGGAAGCATACCATTCAC | FM | |
| Clone 15 | GGGATATTTTATCACCACAG**TGTCAA**TACGGAAGCATACCATTCAC | CQ | |
| Clone 16 | GGGATATTTTATCACCACAG**GTAAAT**TACGGAAGCATACCATTCAC | VN | |
| Clone 17 | GGGATATTTTATCACCACAG**GTGAAC**TACGGAAGCATACCATTCAC | VN | |
| Clone 18 | GGGATATTTTATCACCACAG**GTGAAT**TACGGAAGCATACCATTCAC | VN | |
| Clone 19 | GGGATATTTTATCACCACAG**GTCAAT**TACGGAAGCATACCATTCAC | VN | |
| Clone 20 | GGGATATTTTATCACCACAG**CAAGTA**TACGGAAGCATACCATTCAC | QV | |
| Clone 21 | GGGATATTTTATCACCACAG**ATCTTT**TACGGAAGCATACCATTCAC | IF | |
| Clone 22 | GGGATATTTTATCACCACAG**ATCTTT**TACGGAAGCATACCATTCAC | IF | |
| Clone 23 | GGGATATTTTATCACCACAG**ATTTTC**TACGGAAGCATACCATTCAC | IF | |
| Clone 24 | GGGATATTTTATCACCACAG**TGCGAG**TACGGAAGCATACCATTCAC | CE | |
| Clone 25 | GGGATATTTTATCACCACAG**GTGGAA**TACGGAAGCATACCATTCAC | VE | |
| Clone 26 | GGGATATTTTATCACCACAG**TACAGC**TACGGAAGCATACCATTCAC | YS | |
| Clone 27 | GGGATATTTTATCACCACAG**TATAGC**TACGGAAGCATACCATTCAC | YS | |
| Clone 28 | GGGATATTTTATCACCACAG**TATTCA**TACGGAAGCATACCATTCAC | YS | |
| Clone 29 | GGGATATTTTATCACCACAG**TATTCA**TACGGAAGCATACCATTCAC | YS | |
| Clone 30 | GGGATATTTTATCACCACAG**TATTCA**TACGGAAGCATACCATTCAC | YS | |
| Clone 31 | GGGATATTTTATCACCACAG**GTTGGA**TACGGAAGCATACCATTCAC | **VG** | |
| Clone 32 | GGGATATTTTATCACCACAG**GTTGGA**TACGGAAGCATACCATTCAC | **VG** | |
| Clone 33 | GGGATATTTTATCACCACAG**GTTGGA**TACGGAAGCATACCATTCAC | **VG** | |

### SUPPLEMENTARY MATERIALS

| | | |
|---|---|---|
| Clone 34 | GGGATATTTTATCACCACAG**GCTGGA**TACGGAAGCATACCATTCAC | AG |
| Clone 35 | GGGATATTTTATCACCACAG**TGTGGA**TACGGAAGCATACCATTCAC | CG |
| Clone 36 | GGGATATTTTATCACCACAG**CTTCGG**TACGGAAGCATACCATTCAC | LR |
| Clone 37 | GGGATATTTTATCACCACAG**CTCCGA**TACGGAAGCATACCATTCAC | LR |
| Clone 38 | GGGATATTTTATCACCACAG**CAGCGT**TACGGAAGCATACCATTCAC | QR |
| Clone 39 | GGGATATTTTATCACCACAG**GTCCGA**TACGGAAGCATACCATTCAC | VR |
| Clone 40 | GGGATATTTTATCACCACAG**AATAGG**TACGGAAGCATACCATTCAC | NR |
| Clone 41 | GGGATATTTTATCACCACAG**GGGAGA**TACGGAAGCATACCATTCAC | GR |
| Clone 42 | GGGATATTTTATCACCACAG**AGTCCT**TACGGAAGCATACCATTCAC | SP |
| Clone 43 | GGGATATTTTATCACCACAG**AGTGAT**TACGGAAGCATACCATTCAC | SD |
| Clone 44 | GGGATATTTTATCACCACAG**AAAGGA**TACGGAAGCATACCATTCAC | KG |
| Clone 45 | GGGATATTTTATCACCACAG**AAGTTA**TACGGAAGCATACCATTCAC | KL |
| Clone 46 | GGGATATTTTATCACCACAG**TAGTGG**TACGGAAGCATACCATTCAC | -W |
| Clone 47 | GGGATATTTTATCACCACAG**TAGCGG**TACGGAAGCATACCATTCAC | -R |
| Clone 48 | GGGATATTTTATCACCACAG**TAGGAT**TACGGAAGCATACCATTCAC | -D |
| Clone 49 | GGGATATTTTATCACCACAG**ACTTAA**TACGGAAGCATACCATTCAC | T- |
| Clone 50 | GGGATATTTTATCACCACAG**TCGTAA**TACGGAAGCATACCATTCAC | S- |
| Clone 51 | GGGATATTTTATCACCACAG**CAGTAA**TACGGAAGCATACCATTCAC | Q- |
| Clone 52 | GGGATATTTTATCACCACAG**CGTTGA**TACGGAAGCATACCATTCAC | R- |
| Clone 53 | GGGATATTTTATCACCACAG**TGCTAA**TACGGAAGCATACCATTCAC | C- |
| Clone 54 | GGGATATTTTATCACCACAG**AAATGA**TACGGAAGCATACCATTCAC | K- |
| Clone 55 | GGGATATTTTATCACCACAG**TGATAG**TACGGAAGCATACCATTCAC | -- |

**SUPPLEMENTARY MATERIALS**

Supplementary Table 9: List of oligos used in this study

| Oligo name | Sequence | Purpose |
|---|---|---|
| PexR | CAGGCTTTACACTTTATGCTTCCGGC | Sequencing primers |
| L4440 | AGCGAGTCAGTGAGCGAG | |
| M13F | AGGGTTTTCCCAGTCACGACGTT | |
| P21 | AGGGTTATTGTCTCATGAGCGG | |
| P39 | TACCGCCTTTGAGTGAGCTG | |
| P147 | TTGGGTAACGCCAGGGTTTT | |
| P156 | CGCGTTGGCCGATTCATTAA | |
| pWUS | GACTATACAAAAGTTGGGTAT | |
| A10_OL1 | AGAAGTGAAGCTTGGTCTCATCAGGGGTCATCCAAGAATGTTAT | Clone conversion: MoClo to GreenGate (new overhang is underlined; extra sequences that were added are shown in bold) |
| A10_OL2 | GCCACCATCTGTTCCTTTAACTGCTGAGACCGAATTCTCGCCCT | |
| A12_OL1 | AGAAGTGAAGCTTGGTCTCACTGCGCTCTCAAGATCAAAGGCTT | |
| A12_OL2 | ATTTTAAGATCGCACCATTTACTATGAGACCGAATTCTCGCCCT | |
| C3_OL1 | AGAAGTGAAGCTTGGTCTCAACCTCCAGAAGGTAATTATCCAAG | |
| C3_OL2 | AGAGAAATTTGTAAGTTTGTAACATGAGACCGAATTCTCGCCCT | |
| C7_OL1 | AGAAGTGAAGCTTGGTCTCAGGCT**CCATG**GTGAGCAAGGGCGAGGAGCT | |
| C7_OL2 | AAAAACGCGGCTATTAGATC**A**TCAGTGAGACCGAATTCTCGCCCT | |
| A10_OL1_AS | ATAACATTCTTGGATGACCCCTGATGAGACCAAGCTTCACTTCT | Clone conversion: S/AS experiment (new overhang is underlined) |
| A10_OL2_AS | AGGGCGAGAATTCGGTCTCAGCAGTTAAAGGAACAGATGGTGGC | |
| A12_OL1_AS | AAGCCTTTGATCTTGAGAGCGCAGTGAGACCAAGCTTCACTTCT | |
| A12_OL2_AS | AGGGCGAGAATTCGGTCTCATAGTAAATGGTGCGATCTTAAAAT | |
| GST_FW | ATGTCCCCTATACTAGGTTA | One PCR to many entries (new overhang is underlined; extra sequences that were added are shown in bold) |
| GST_REV | ACGCGGAACTAGATCCGATT | |
| A-GST-B_L | AGAAGTGAAGCTTGGTCTCAACCTATGTCCCCTATACTAGGTTA | |
| A-GST-B_R | AATCGGATCTAGTTCCGCGT**TC**AACATGAGACCGAATTCTCGCCCT | |
| B-GST-C_L | AGAAGTGAAGCTTGGTCTCAAACAATGTCCCCTATACTAGGTTA | |
| B-GST-C_R | AATCGGATCTAGTTCCGCGTGGCTTGAGACCGAATTCTCGCCCT | |
| C-GST-D_L | AGAAGTGAAGCTTGGTCTCAGGCT**CT**ATGTCCCCTATACTAGGTTA | |
| C-GST-D_R | AATCGGATCTAGTTCCGCGTTCAGTGAGACCGAATTCTCGCCCT | |
| D-GST-E_L | AGAAGTGAAGCTTGGTCTCATCAG**CT**ATGTCCCCTATACTAGGTTA | |
| D-GST-E_R | AATCGGATCTAGTTCCGCGTCTGCTGAGACCGAATTCTCGCCCT | |

**SUPPLEMENTARY MATERIALS**

| | | |
|---|---|---|
| E-GST-F_L | AGAAGTGAAGCTTGGTCTCACTGC**CT**ATGTCCCCTATACTAGGTTA | |
| E-GST-F_R | AATCGGATCTAGTTCCGCGT**TGA**ACTATGAGACCGAATTCTCGCCCT | |
| 3xHA_FW | ATGGCATACCCTTACGATGT | |
| 3xHA_REV | AGCGTAATCTGGAACGTCG | |
| A-3xHA-B_L | AGAAGTGAAGCTTGGTCTCAACCTATGGCATACCCTTACGATGT | |
| A-3xHA-B_R | ACGACGTTCCAGATTACGCT**TC**AACATGAGACCGAATTCTCGCCCT | |
| B-3xHA-C_L | AGAAGTGAAGCTTGGTCTCAAACAATGGCATACCCTTACGATGT | |
| B-3xHA-C_R | ACGACGTTCCAGATTACGCTGGCTTGAGACCGAATTCTCGCCCT | |
| C-3xHA-D_L | AGAAGTGAAGCTTGGTCTCAGGCT**CT**ATGGCATACCCTTACGATGT | |
| C-3xHA-D_R | ACGACGTTCCAGATTACGCTTCAGTGAGACCGAATTCTCGCCCT | |
| D-3xHA-E_L | AGAAGTGAAGCTTGGTCTCATCAG**CT**ATGGCATACCCTTACGATGT | |
| D-3xHA-E_R | ACGACGTTCCAGATTACGCTCTGCTGAGACCGAATTCTCGCCCT | |
| E-3xHA-F_L | AGAAGTGAAGCTTGGTCTCACTGC**CT**ATGGCATACCCTTACGATGT | |
| B_GFP_C_L | AGAAGTGAAGCTTGGTCTCAAACACTATGGTGAGCAAGGGCGAG | Clone conversion: GreenGate to GreenGate (new overhang is underlined; extra sequences that were added are shown in bold) |
| B_GFP_C_R | GCATGGACGAGCTGTACAAGGGCTTGAGACCGAATTCTCGCCCT | |
| B_MBP_C_L | AGAAGTGAAGCTTGGTCTCAAACAGCATGAAAATCGAAGAAGGTAAACTG | |
| B_MBP_C_R | CCCTGAAAGACGCGCAGACTGGCTTGAGACCGAATTCTCGCCCT | |
| B_MtU6_C_L | AGAAGTGAAGCTTGGTCTCAAACAATGCCTATCTTATATGATCA | |
| B_MtU6_C_R | CTTGTACAAAGTTGGCATTAGGCTTGAGACCGAATTCTCGCCCT | |
| C_MtU6_D_L | AGAAGTGAAGCTTGGTCTCAGGCTATGCCTATCTTATATGATCA | |
| C_MtU6_D_R | CTTGTACAAAGTTGGCATTATCAGTGAGACCGAATTCTCGCCCT | |
| A-GST-B_L | AGAAGTGAAGCTTGGTCTCAACCTATGTCCCCTATACTAGGTTA | |
| A-GST-B_R | AATCGGATCTAGTTCCGCGT**TC**AACATGAGACCGAATTCTCGCCCT | |
| A-MBP-B_L | AGAAGTGAAGCTTGGTCTCAACCTATGAAAATCGAAGAAGGTAAACT | |
| A-MBP-B_R | CCCTGAAAGACGCGCAGACT**TC**AACATGAGACCGAATTCTCGCCCT | |
| A10_AarI_OL1 | ACACTATAGAAGTGAAGCTT**CACCTGC**AATATCAGGGGTCATCCAAGAATGTTAT | Replacement of Type IIS recognition site (new Type IIS recognition site in bold; new overhang is underlined) |
| A10_AarI_OL2 | GCCACCATCTGTTCCTTTAACTGCTCGT**GCAGGTG**GAATTCTCGCCCTATAGTGA | |
| A12_SapI_OL1 | ACACTATAGAAGTGAAGCTT**GCTCTTC**ATGAGCTCTCAAGATCAAAGGCTT | |
| A12_SapI_OL2 | ATTTTAAGATCGCACCATTTGATT**GAAGAGC**GAATTCTCGCCCTATAGTGA | |
| GFP_StitAmp_F | ATGGTGAGCAAGGGCGAGG | GFP amplification |
| GFP_StitAmp_R | CTTGTACAGCTCGTCCATGCC | |
| GUS_GFP_OL1 | AGAAGTGAAGCTTGGTCTCAGGCTCAACAATGGTCCGTCCTGTA | |

**SUPPLEMENTARY MATERIALS**

| | | |
|---|---|---|
| GUS_GFP_OL2 | CGCAGCAGGGAGGCAAACAA**GGTGGATCAGGCGGAAGT**ATGGTGAGCAAGGGCGAGGA | Clone tagging with PCR products or donor plasmids (overhang sequence is underlined; extra sequence that was added is in bold) |
| GUS_GFP_OL3 | GCATGGACGAGCTGTACAAGTCAGTGAGACCGAATTCTCGCCCT | |
| INT4_V5_OL1 | AGAAGTGAAGCTTGGTCTCAGGCTCCATGATTACGACCAGAAAG | |
| INT4_V5_OL2 | TAGATATACATTGGACCTTTGGTAAGCCAATCCCTAATCC | |
| INT4_V5_OL3 | TCGGACTCGACTCAACCTAATCAGTGAGACCGAATTCTCGCCCT | |
| Csy4_mCherry_OL1 | AGAAGTGAAGCTTGGTCTCAAACAATGGTGAGCAAGGGCGAGGA | |
| Csy4_mCherry_OL2 | GCATGGACGAGCTGTACAAG**GGAGGTTCA**ATGGATCATTATCTTGATAT | |
| Csy4_mCherry_OL3 | TTGAAGAAAATCCTGGACCCGGCTTGAGACCGAATTCTCGCCCT | |
| INT4_left | AGAAGTGAAGCTTGGTCTCAGGCTCCATGATTACGACCAGAAAG | Clone tagging with oligo-encoded sequences (overhang sequence is underlined; extra sequence that was added is in bold) |
| INT4_ER_left | AGAAGTGAAGCTTGGTCTCAGGCTCCATG**AAGGTACAGGAGGGTTTGTTCGTGGTGGCTGTTTTCTACCTTGCTTATACGCAGCTAGTCAAGGGG**ATGATTACGACCAGAAAG | |
| INT4_SV40_right | TAGATATACATTGGACCTTT**CCTAAGAAGAAGAGGAAGGTT**TCAGTGAGACCGAATTCTCGCCCT | |
| INT4_right | TAGATATACATTGGACCTTTTCAGTGAGACCGAATTCTCGCCCT | |
| FT_Oligo | TTGGCCATAAGTAACCTTTAGAGTGATTGATCTATTAAACGGATCAAGAACGTCTCCAACAACTCTGCTTACTATAAGAGGGTCTCTTATATTTATAGACATGCTTCTTGGTGCCGCGCCT | pegRNA assembly |
| gRNA_Spacer_FT | TAAAGGTTACTTATGGCCAACTCGTGACCACCTTCACCCAGTTTTAGAGCTAGAAATAGC | |
| gRNA-Scaffold | GCACCGACTCGGTGCCACTTTTTCAAGTTGATAACGGACTAGCCTTATTTTAACTTGCTATTTCTAGCTCTAAAAC | |
| gRNA_RTPBS_TEVOPREQ1 | AAGTGGCACCGAGTCGGTGCTGCACGCCGTACGTGAAGGTGGTCACCGCGGTTCTATCTAGTTACG | |
| TEVOPREQ1_Oligo | TCGGTCTCAATACAAAAAAATTCTAGTTGGTTTAACGCGTAACTAGATAGAACCGCG | |
| gRNA_Spacer | AGGCGCGGCACCAAGAAGCACTCGTGACCACCTTCACCCAGTTTTAGAGCTAGAAATAGC | |
| gRNA_RTPBS | AAGTGGCACCGAGTCGGTGCTGCACGCCGTACGTGAAGGTGGTCACTTTTTTTGTATTGAGACCGA | |
| OLE1P_XbaI_OL1 | TATCCATTTTCTTCATTGT**TATAGA**ATGTCGCGGAACAAATTTT | Restriction enzyme recognition site removal (The modified restriction enzyme motif is shown in bold, with the changed base underlined. The original sequence is "TCTAGA" for XbaI, "GAATTC" for EcorI, |
| OLE1P_XbaI_OL2 | CGATCTGATACTGATAACG**TCTCGA**TTTTTAGGGTTAAAGCAAT | |
| Csy4_EcorI_OL1 | AGACTTGGAGAAAGACTTA**GGATTC**ATGCTTCTGCTGATGATCT | |
| Csy4_EcorI_OL2 | GAAGAAGAAGCTAGAAAAA**GAGTTC**CTGATACTGTTGCTAGAGC | |
| Csy4_EcorI_OL3 | CTGGACCCGGCTTGAGACC**GAACTC**TCGCCCTATAGTGAGTCGT | |
| tUM140_1 | AGAAGTGAAGCTTGGTCTCACTGCGCCGCGCACAGCTGACGTAG | |

**SUPPLEMENTARY MATERIALS**

| | | |
|---|---|---|
| tUM140_2a | AACTCTAGTGTAATCCCAGACTGGTG**T**GACCAGTGACATTGACACCA | “GAGACC” for BsaI, “CACCTGC“ for PaqCI, and “GAAGAC” for BbsI). |
| tUM140_2b | AACTCTAGTGTAATCCCAGACTGGTGAG**G**CCAGTGACATTGACACCA | |
| tUM140_2c | AACTCTAGTGTAATCCCAGACTGGTGA**C**ACCAGTGACATTGACACCA | |
| tUM140_2d | AACTCTAGTGTAATCCCAGACTGGTGAGA**A**CAGTGACATTGACACCA | |
| tUM140_2e | AACTCTAGTGTAATCCCAGACTGGT**T**AGACCAGTGACATTGACACCA | |
| tUM140_3 | TACATTTGCATGCAAAGTTGACTATGAGACCGAATTCTCGCCCT | |
| MoClo_mCherry_1 | TTAATCACTCTGTGGTCTCATTCGATGGTGAGCAAGGGCGAGGA | |
| Alt_mCherry_1 | AGAAGTGAAGCTTGGTCTCATTCGCTATGGTGAGCAAGGGCGAGGA | |
| MoClo_mCherry_2 | GGCCCCGTAATGCAGAAGAA**A**ACCATGGGCTGGGAGGCCTCCTCCGAGC | |
| MoClo_mCherry_3 | TGGACGAGCTGTACAAGTAGGCTTTGAGACCACGAAGTGGCTCT | |
| Alt_mCherry_3 | GCATGGACGAGCTGTACAAGGCTTTGAGACCGAATTCTCGCCCT | |
| MoClo_WUS_1 | TTAATCACTCTGTGGTCTCAGGAGCCCATGTTTACGTTTACGTT | |
| Alt_WUS_1 | AGAAGTGAAGCTTGGTCTCAGGAGCCCATGTTTACGTTTACGTT | |
| MoClo_WUS_2 | TACTCAACATGTTCATAAGTACACCTG**A**CTTCACACTCGTTTCACACA | |
| MoClo_WUS_3 | CAAAAGTCGAATCAAACACACCCATTGAGACCACGAAGTGGCTCT | |
| Alt_WUS_3 | CAAAAGTCGAATCAAACACACCCATTGAGACCGAATTCTCGCCCT | |
| GUS_Aar1 | AAGACTGTAACCACGCGTCTGTTGACTGGCA**A**GTGGTGGCCAATGGTGATGT | |
| GUS_Aar1b | AAGACTGTAACCACGCGTCTGTTGACTGGCAGGT**C**GTGGCCAATGGTGATGTCAG | |
| GUS_Aar2 | CCGACGCGTCCGATCACCTG**T**GTCAATGTAATGTTCTGCGACGCTCACA | |
| IncLib_Oligo1 | TCCAAGCTCAAGCTAAGCTTACCTCCAGAAGGTAATTATCCAAG | Stitching oligos for incompatible library assembly |
| IncLib_Oligo2 | AGAGAAATTTGTAAGTTTGTATGCCATGGTGAGCAAGGGCGAG | |
| IncLib_Oligo3 | GCATGGACGAGCTGTACAAGCTAACCCCGATGAGCTAAGCTAGC | |
| IncLib_Oligo4 | CATGTACTCGACGGCCGAGTGTATAAGCTTACCTTACTTAGATC | |
| amilCPOrange_Mut Oligo | GGGATATTTTATCACCACAG**NNNNNN**TACGGAAGCATACCATTCAC | Saturation mutagenesis of amilCP Orange (ambiguous bases are shown in bold) |
| Orange_Seq_F | AATAGGCGTATCACGAGGC | Colony PCR primers for saturation mutagenesis experiment |
| Orange_Seq_R | AGCGAGTCAGTGAGCGAG | |
| Orange_NGS_F | GGAGCAGACGGTAAAGCTCA | Primers for generation of NGS amplicon of amilCP_Orange |
| Orange_NGS_R | AGTTGCCTTGGATGCTGGAA | |
