## Supplementary Figure for "Simple and efficient modification of Golden Gate design standards and parts using oligo stitching"

### SUPPLEMENTARY FIGURES

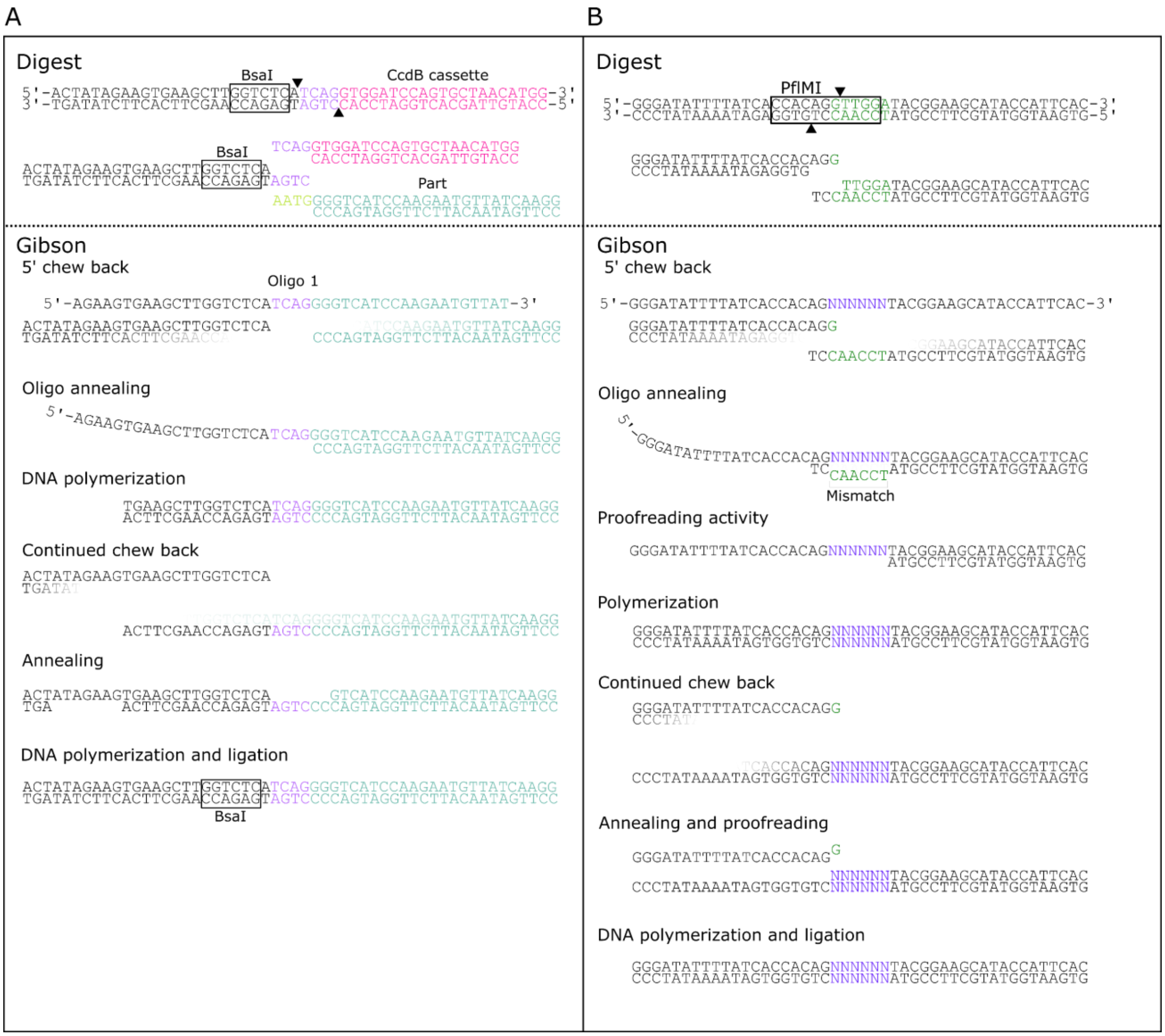


**Supplementary Figure 1. Mechanism of Gibson assembly using oligos.** (A) Oligo stitching mechanism to change a Type IIS restriction enzyme generated overhangs. Only the 5' side of the reaction is shown for simplicity. The acceptor vector containing a CcdB-cassette and the donor vector (backbone is omitted) are digested by BsaI. The digested vectors are mixed with two oligos that have homology to both the backbone (20 bp in black) and the part (20 bp in teal). Between these regions is the 4-bp overhang (purple). In the Gibson assembly reaction, the 5' ends will be chewed back by the 5' DNA exonuclease. The oligos will then anneal to the complimentary sequences and used as a template by DNA polymerase. The exonuclease continues to chew back the 5' ends, both of the newly synthesized sequence, as well as of the backbone. These two sequences can then anneal, polymerize and be ligated together to form a new double stranded flank containing the new overhang. (B) Mechanism for reactions where 3' cheback from the proofreading activity of the DNA polymerase is needed. In this example the reaction is shown that was used for the generation of the saturation mutagenesis screen of amilCP_Orange. The same reactions take place as in A, but because PflMI cuts upstream of where the modifications are needed, a 3' chew back is needed.

### SUPPLEMENTARY FIGURES

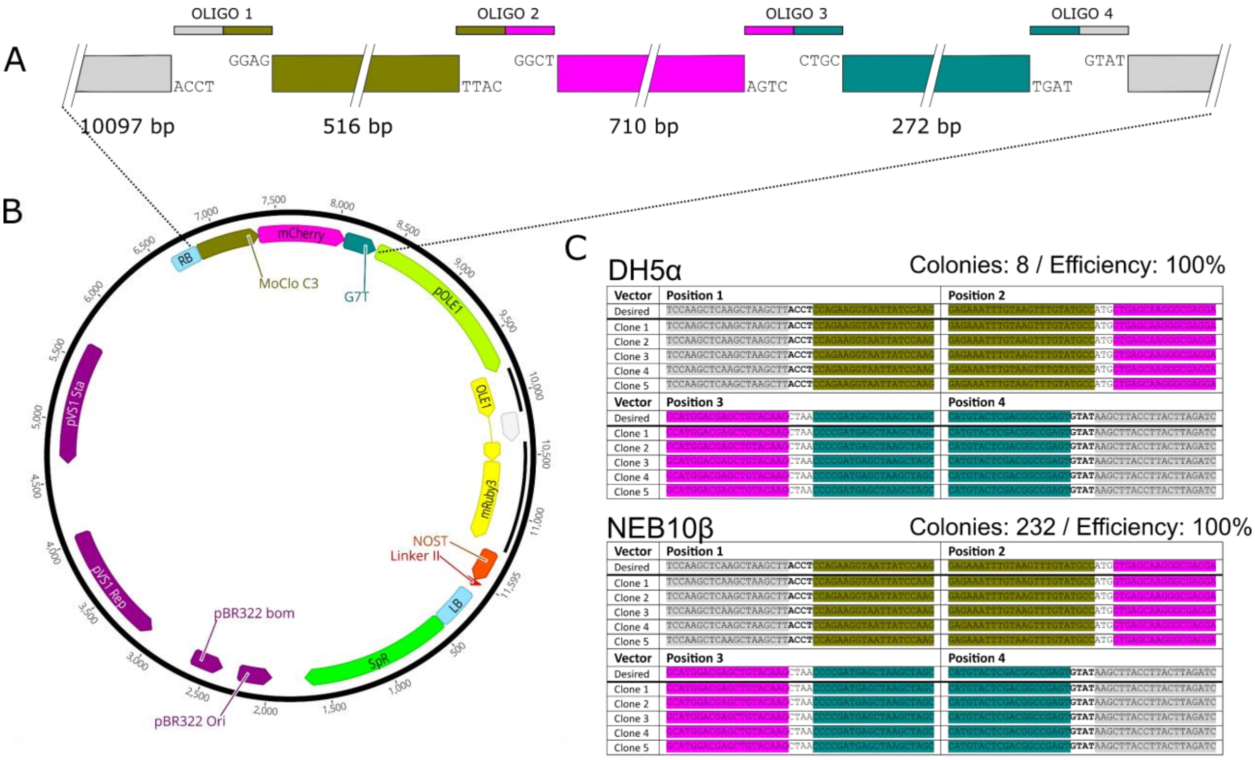


**Supplementary Figure 2. Incompatible library assembly.** (A) The different incompatible parts assembled in this experiment. The circular backbone (10,097 bp) is depicted in gray, the Cassava Vein Mosaic Virus promoter (516 bp) in brown, the mCherry CDS (710 bp) in pink and the G7T terminator (272 bp) in teal. Note that none of the overhangs are compatible for Golden Gate assembly. (B) Fully assembled 11,595 bp construct. (C) Sequence comparison of desired sequence at the four junctions and the sequences of 5 randomly picked clones. Both DH5α chemically competent cells and NEB10B electrocompetent cells were used. The number of colonies and the cloning efficiency is also indicated.

### SUPPLEMENTARY FIGURES

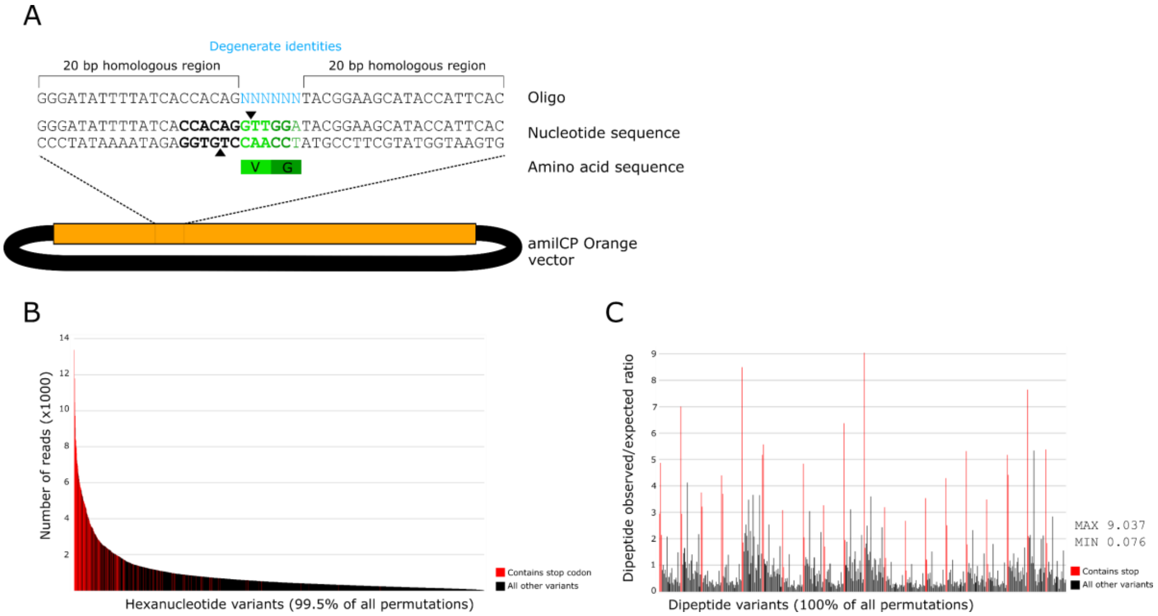


**Supplementary Figure 3. Supporting information for saturation mutagenesis experiment.** (A) Schematic overview of the amilCP_Orange chromoprotein vector showing the nucleotides centered around the codons V63 and G64. The recognition site of the restriction enzyme PflMI is indicated in bold and its cutting position by triangles. The mutagenic oligo is also indicated and was composed out of 20 nt homology on either end of the six degenerate nucleotides. (B) Distribution of the hexanucleotide variants present in the reads and ranked according to abundance. Data shown in red indicates that the hexanucleotide encoded at least one stop codon in either of the two codons. The original variant is not included in the graph. (C) Ratio of observed and expected of each of the possible 441 dipeptides/amino acid-stop codon/stop codon-stop codon combinations. Data shown in red indicate that at least one stop codon was present. The maximum and minimum observed/expected ratio is also indicated. The original codon combination of the original variant is not included in the analysis.
