## Supplementary protocol for "Simple and efficient modification of Golden Gate design standards and parts using oligo stitching"

**The theory**

Oligos can be used to modify Golden Gate parts in various ways with Gibson assembly in a process called "oligo stitching". Oligo stitching can be used to change the overhang sequence between standards (e.g. MoClo to GreenGate) and also within a standard. Oligos are designed to have 20 bp homology to the backbone and 20 bp to the part, with 4 bp in the middle specifying the new overhang and/or other elements (tags, start/stop codons, etc.). The orientation of the oligos does not appear to have an effect.

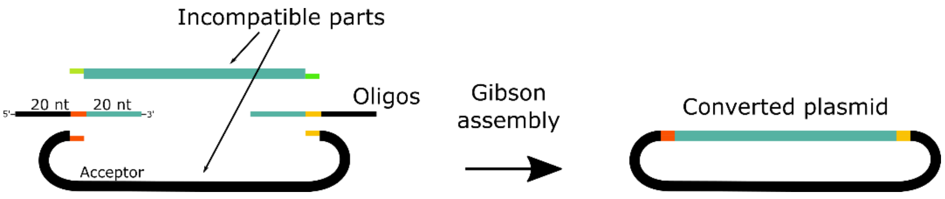


Figure 1. Oligo stitching to change overhangs on Golden Gate Plasmid.

Oligo stitching can also be used to assemble non-compatible parts allowing one to mix and match different Golden Gate standards.

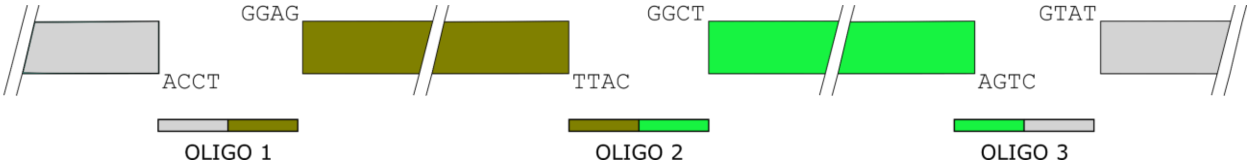


**Oligos can be directly assembled** into a synthetic DNA sequence. For sequences of ≥300 bp, DNA synthesis is probably a more economic option, but for sequences composed of a combination of standard (fixed) parts and variable parts, this is a convenient solution. For example, prime editing gRNAs are composed of a variable 60 bp spacer sequence (20 bp backbone homology--20 bp spacer--20 bp scaffold homology), a fixed scaffold and variable RT+PBS (20 bp scaffold homology--X bp of RT+PBS--20 bp of backbone homology). As oligos longer than 60 bp are significantly more expensive and take more time to arrive, the homologies can be shortened (e.g., 17 bp on both sides instead of 20 bp). This will reduce efficiency but still yield sufficient colonies. Note that we have only performed these assemblies swapping sense and antisense orientations. The design is based on Kweon et al., 2021.

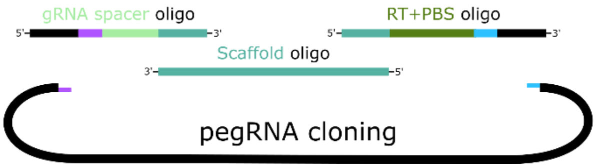


**Plasmids can be mutagenized** to remove an unwanted restriction site (domestication) or perform saturation mutagenesis. Note that success depends on the restriction enzyme used (BsaI led to more contaminating original plasmid than with XbaI and EcoRI). Alternatively, by using oligos containing ambiguous bases a library of plasmid variants can be produced. For mutating one codon for example, take 20 bp upstream of the codon, add 3 ambiguous bases, and then add 20 bp downstream of the codon for the oligo design. Multiple mutagenic oligos can also be used in one reaction.

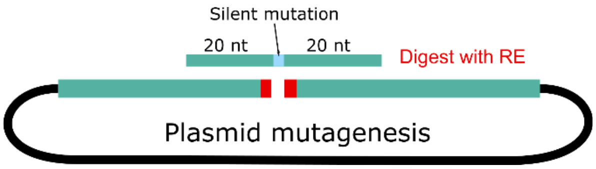

**Supplementary protocol**

**Step-by-step protocol**

**Oligos:** Resuspend the oligos to 100 µM and take 1 µL (100 pmol) of each and add to 331 µL of water. Per oligo concentration 0.3 pmol/ul = 300 fmol.

For ≥2 oligos, add 1 µL per *n* oligo part to 333-*n* µL of water.

For DNA assembly: Dilute oligos to 100 nM. Add 1 µL (100 fmol) of each oligo to Gibson assembly reaction below. Note: For 3-4 oligos, $10^6$ competent cells can be used, for >4 oligos higher efficiencies are recommended.

**Plasmids and dsDNA parts** may be prepared by digest or PCR.

| Component | Amount |
|---|---|
| Part Plasmid | 1 µg |
| 10X Buffer* | 2 µL |
| Enzyme | 0.5 µL |
| $H_2O$ | variable |
| Total | 20 µL |

| Component | Amount |
|---|---|
| Acceptor Plasmid | 1 µg |
| 10X Buffer* | 2 µL |
| Enzyme | 0.5 µL |
| $H_2O$ | variable |
| Total | 20 µL |

*Don't forget to add Bovine Serum Albumin (BSA) if using Promega enzymes

Digest for ~4 hours.

Heat inactivate according to the manufacturer's instructions, typically 60 °C or 80°C for 20 minutes.

Prepare PCR parts as any other standard cloning reaction.

PCR parts may be used unpurified or may require gel purification if multiple bands are present. Gel extract plasmid parts with the same resistance marker to avoid picking up the original vector. Alternatively, we have GreenGate entry vectors containing Gm and TetR resistance, as well as SpecR for non-GreenGate.

**Gibson Assembly**

Determine the ng of plasmid needed for 0.04 pmol of parts.

$$ng\ of\ plasmid\ needed = bp\ of\ plasmid\ \times 0.0264$$

Use the graph at the left to quickly estimate →

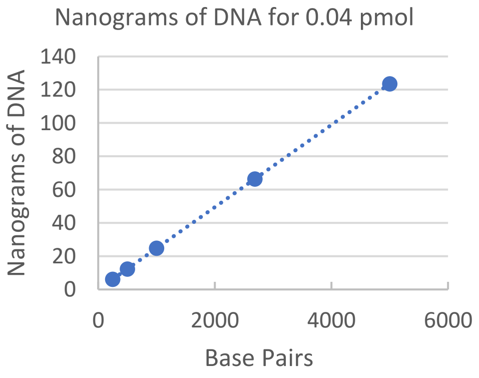


Prepare the NEBuilder reaction as follows

| Component | Amount | Volume (µL) |
|---|---|---|
| Oligo mix | 0.30 pmol | 1 |
| Part 1 | 0.04 pmol | variable |
| Part 2… | 0.04 pmol | variable |
| Acceptor | 0.04 pmol | variable |
| 2XNEBuilder | | 5 |
| $H_2O$ | | variable |
| Total | | 10 µL |

Incubate at 50°C for one hour and transform ~2 µl into ~25 µl *E. coli*. Reactions can be scaled up or down as necessary.

Notes: NEBuilder is essential for reactions that chew back 3' ends. 1 µL of each digested plasmid (~50 ng) will also yield satisfactory results. We obtained good results with up to three parts, but cells with higher transformation efficiency (>$10^6$) may be needed when more parts are used.

**Quality Control:** Sanger sequencing. Errors may occur at the junctions, so clones should be sequenced at all junctions.
